## Supplementary Information for "Spatial redundancy transformer for self-supervised fluorescence image denoising"

### CONTENT

#### I. Supplementary Figures


| | |
|---|---|
| **Supplementary Figure 1** | Sampling strategies of different self-supervised denoising methods. |
| **Supplementary Figure 2** | Denoising performance of different neighbor sampling schemes. |
| **Supplementary Figure 3** | Evaluating the data dependency of SRDTrans. |
| **Supplementary Figure 4** | Examples of simulated calcium imaging data with different SNRs. |
| **Supplementary Figure 5** | Fourier spectrum of SRDCNN and SRDTrans. |
| **Supplementary Figure 6** | Performance of SRDCNN and SRDTrans with different input temporal scales (30-Hz calcium imaging data). |
| **Supplementary Figure 7** | Performance of SRDCNN and SRDTrans with different input temporal scales (1-Hz calcium imaging data). |
| **Supplementary Figure 8** | Investigating the generalization ability of SRDTrans by cross-dataset and cross-modality validation. |
| **Supplementary Figure 9** | Examples of simulated SMLM data with different SNRs. |
| **Supplementary Figure 10** | Performance of various self-supervised methods. |
| **Supplementary Figure 11** | Resolution analysis of reconstructed SMLM images. |
| **Supplementary Figure 12** | Localization accuracy before and after denoising. |
| **Supplementary Figure 13** | Spatiotemporal analysis of denoised calcium imaging data. |
| **Supplementary Figure 14** | Frequency domain analysis of different denoising methods. |
| **Supplementary Figure 15** | Denoising experimentally obtained calcium imaging data with synchronized high-SNR (10-fold photons) reference. |
| **Supplementary Figure 16** | Comparing the performance of DeepInterpolation, DeepCAD, and SRDTrans. |

| | |
|---|---|
| **Supplementary Figure 17** | Performance of DeepCAD and SRDTrans at different imaging speeds. |
| **Supplementary Figure 18** | Pearson correlations at different imaging speeds. |
| **Supplementary Figure 19** | Comparing DeepCAD and SRDTrans on moving objects with different moving speeds. |
| **Supplementary Figure 20** | The applicability of SRDTrans on confocal microscopy and light sheet microscopy. |
| **Supplementary Figure 21** | The mechanism of feature extraction in 3D-UNet and SRDTrans. |
| **Supplementary Figure 22** | Architecture of the spatiotemporal transformer block (STB). |
| **Supplementary Figure 23** | Convergence curves of different loss functions. |
| **Supplementary Figure 24** | Training stability of SRDTrans. |
| **Supplementary Figure 25** | Comparing the denoising performance of SRDTrans on Gaussian and Poisson noise. |


### II. Supplementary Tables


| | |
|---|---|
| **Supplementary Table 1** | Model complexity of CNN and SRDTrans. |
| **Supplementary Table 2** | Model complexity and denoising performance of different transformer-style networks. |
| **Supplementary Table 3** | Quantitative comparison of SRDCNN, DeepCAD, and SRDTrans at different imaging speeds. |
| **Supplementary Table 4** | Parameters for the SMLM experiments. |
| **Supplementary Table 5** | Training configurations of different self-supervised denoising methods. |


### III. Supplementary Notes


| | |
|---|---|
| **Supplementary Note 1** | Proof of spatial redundancy sampling scheme. |
| **Supplementary Note 2** | The spectral bias of CNNs. |

### IV. Supplementary Videos

| | |
|---|---|
| **Supplementary Video 1** | The denoising performance of different methods on calcium imaging data sampled at 0.3 Hz. |
| **Supplementary Video 2** | SRDTrans massively improves the SNR of SMLM. |
| **Supplementary Video 3** | SRDTrans massively improves the SNR of calcium imaging. |
| **Supplementary Video 4** | Comparing the performance of DeepCAD and SRDTrans on fast-moving objects. |
| **Supplementary Video 5** | SRDTrans enhances two-photon volumetric calcium imaging in the mouse cortex. |

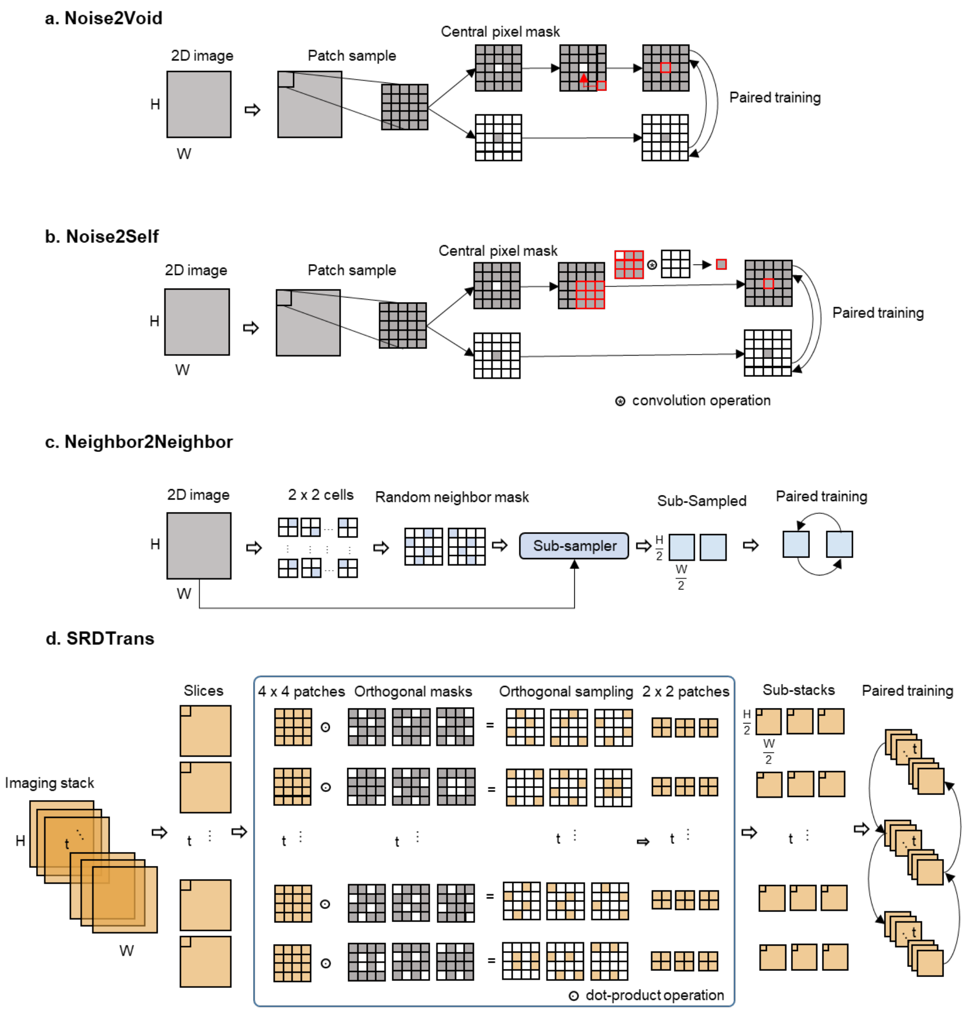


#### Supplementary Figure 1

**Sampling strategies of different self-supervised denoising methods.**

Comprehensive demonstration of different sampling strategies, including Noise2Void[1] (**a**), Noise2Self[2] (**b**), Neighbor2Neighbor[3] (**c**) and SRDTrans (**d**). **a-b**, Both Noise2Void and Noise2Self employ the "blind-spot" strategy. The raw image is randomly split into a series of patches and for each patch a randomly selected pixel is masked. In Noise2Void, the masked pixel is replaced with a neighbor pixel; while in Noise2Self, this pixel is replaced with the convolution of a hollow 3×3 window. Then the original patch and the masked patch are used for paired training. **c**, In Neighbor2Neighbor, neighboring sub-sampling is applied to generate the two sub-sampled images for paired training. In each 2×2 cell of the raw images, two neighboring pixels are randomly selected, and rearranged into two sub-images, respectively. **d**, In SRDTrans, the image stack (xy-t or xy-z) was sampled with three orthogonal masks to generate three sub-stacks. The central sub-stack is selected as the training input, and its two adjacent sub-stacks are designated as the corresponding training targets. The sampling strategy guarantees the pixels of each input-target pair are vertically or horizontally adjacent.

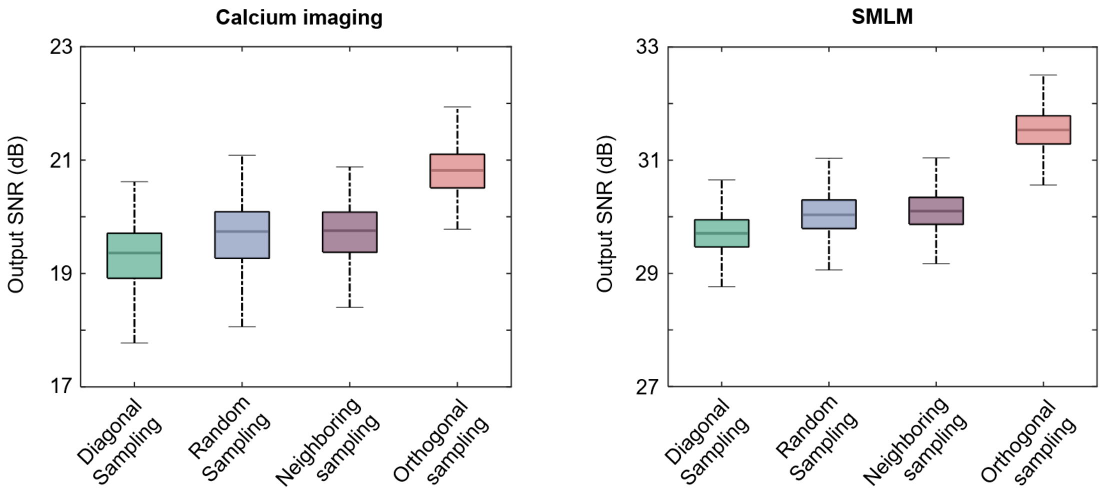


**Supplementary Figure 2**

**Denoising performance of different neighbor sampling schemes.**

We quantitatively compared the performance of different neighbor sampling schemes both on simulated calcium imaging (1000 frames, 1 Hz frame rate, input SNR=-0.86 dB) and SMLM (6000 frames, 200 Hz frame rate, input SNR=9.76 dB) data. Four sampling schemes were investigated, including (1) diagonal sampling that only randomly selects diagonal pixels, (2) random sampling that selects horizontally or vertically or diagonally adjacent pixels, (3) neighboring sampling that selects horizontally or vertically adjacent pixels, (4) orthogonal sampling that simultaneously selects horizontally and vertically adjacent pixels. Left, statistical results on calcium imaging data (N=1000). Right, statistical results on SMLM data (N=6000). Among them, diagonal sampling has the worst performance because diagonal pixels have the longest distance in a 2×2 patch and their similarity is the lowest. The proposed orthogonal sampling has the best performance since it can avoid using diagonal pixels and learn horizontal and vertical correlations isotropically.

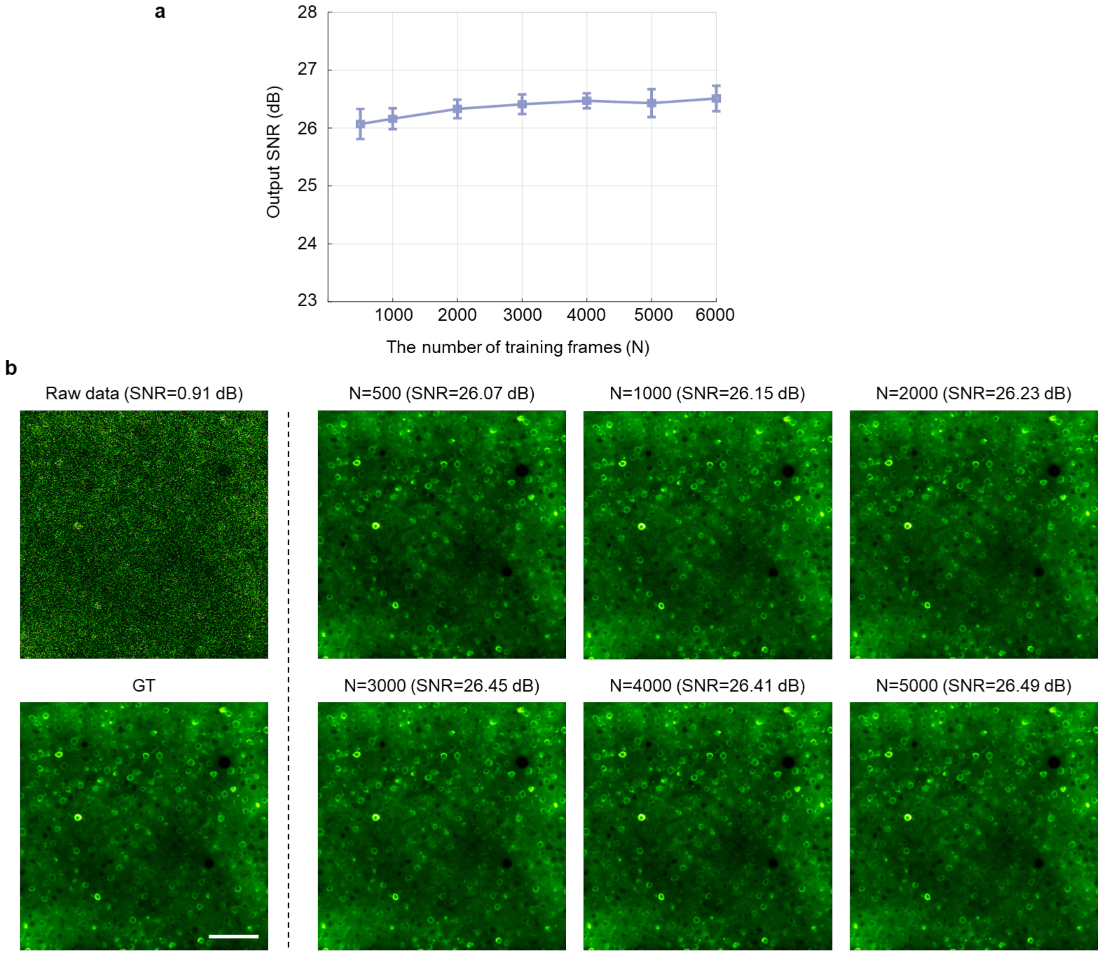


#### Supplementary Figure 3

**Evaluating the data dependency of SRDTrans.**

We trained SRDTrans models with simulated calcium imaging data (30 Hz frame rate, SNR=0.91 dB). The size of each image stack is 490×490×N, where N denotes the number of frames from {500, 1000, 2000, 3000, 4000, 5000, 6000}. Each model was trained for 20 epochs. **a**, Quantitative evaluation of the denoising performance trained with different amounts of data (N). The lines connect mean values and the error bars represent the standard deviation. The sample size is the number of training frames (N). **b**, Representative denoising results of models trained with different amounts of data. The raw noisy data and corresponding ground truth (GT) are presented in the leftmost panel. Scale bar, 100 μm.

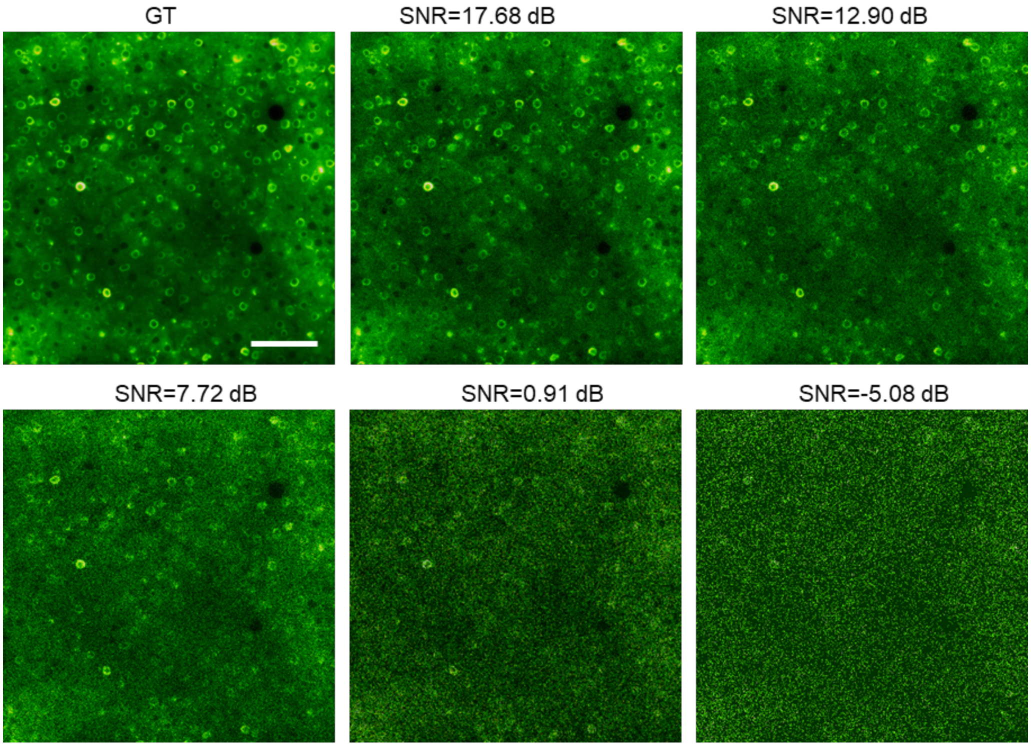


**Supplementary Figure 4**

**Examples of simulated calcium imaging data with different SNRs.**

We used Neural Anatomy and Optical Microscopy (NAOMi)[4] to simulate two-photon calcium imaging data. We first generated the noise-free time-lapse image sequence containing high-fidelity neuronal structures and dynamics, and then added Mixed Poisson Gaussian (MPG) noise to synthesize authentic low-SNR images. Representative images at different SNR levels corresponding to the same ground truth (GT) are presented. Scale bar, 100 µm.

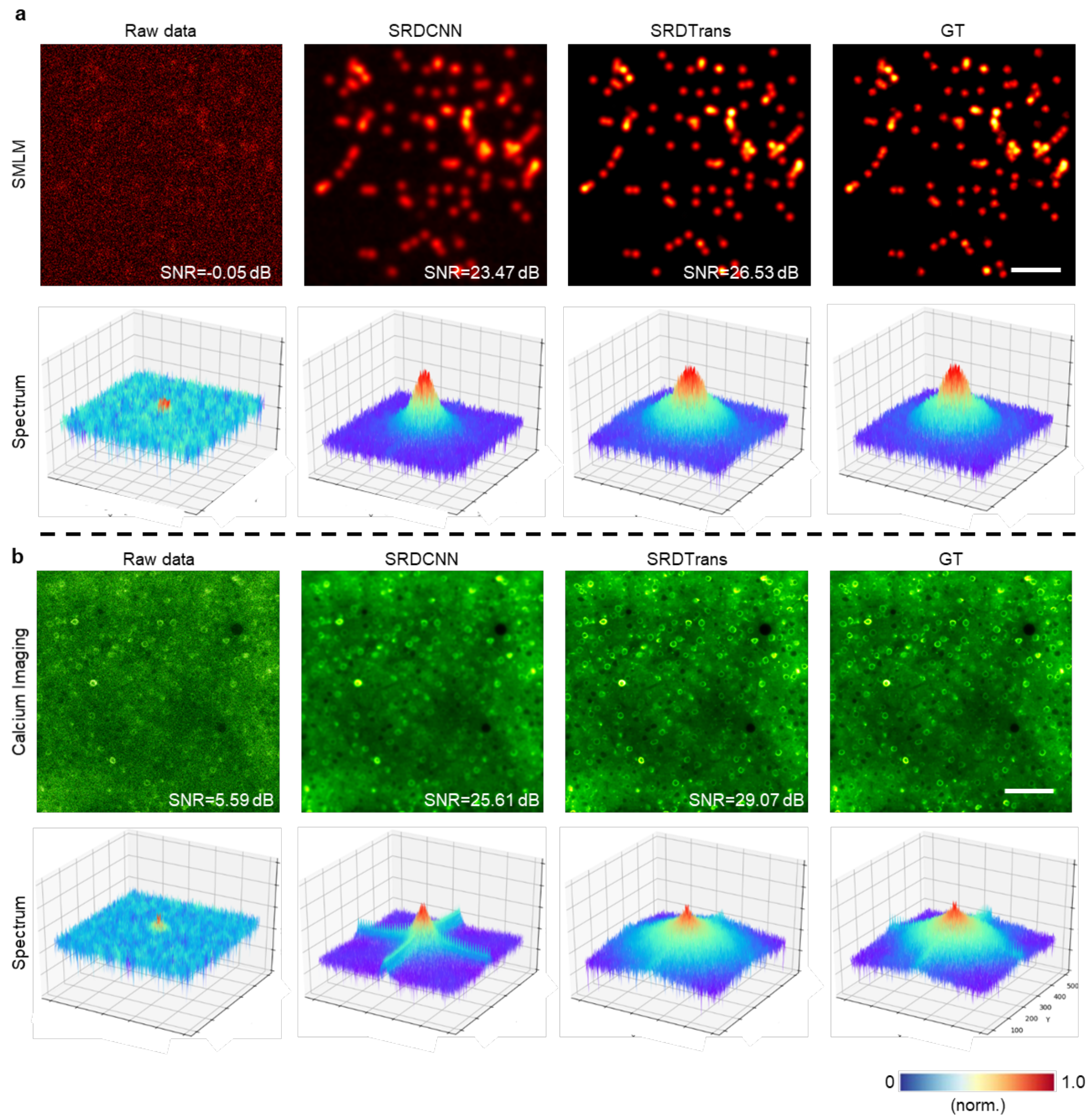


**Supplementary Figure 5**

**Fourier spectrum of SRDCNN and SRDTrans.**

Spatial images and corresponding Fourier spectrum of SMLM (**a**) and calcium imaging (**b**). The raw noisy data, SRDCNN denoised data, SRDTrans denoised data, and ground truth (GT) are presented. Simulated SMLM (24000 frames, 200 Hz frame rate, SNR=-0.05 dB) and calcium imaging data (6000 frames, 30 Hz frame rate, SNR=5.59 dB) were used to train specific models for each method. The Fourier spectrum was calculated by applying discrete Fourier transform to the spatial image. In the raw noisy data, frequency information is nearly overwhelmed by noise. Limited by spectral bias[5-8], SRDCNN can only recover low-frequency information. In contrast, SRDTrans can recover most of the high-frequency information and is highly consistent with GT. Scale bar, 2 μm in **a** and 100 μm in **b**.

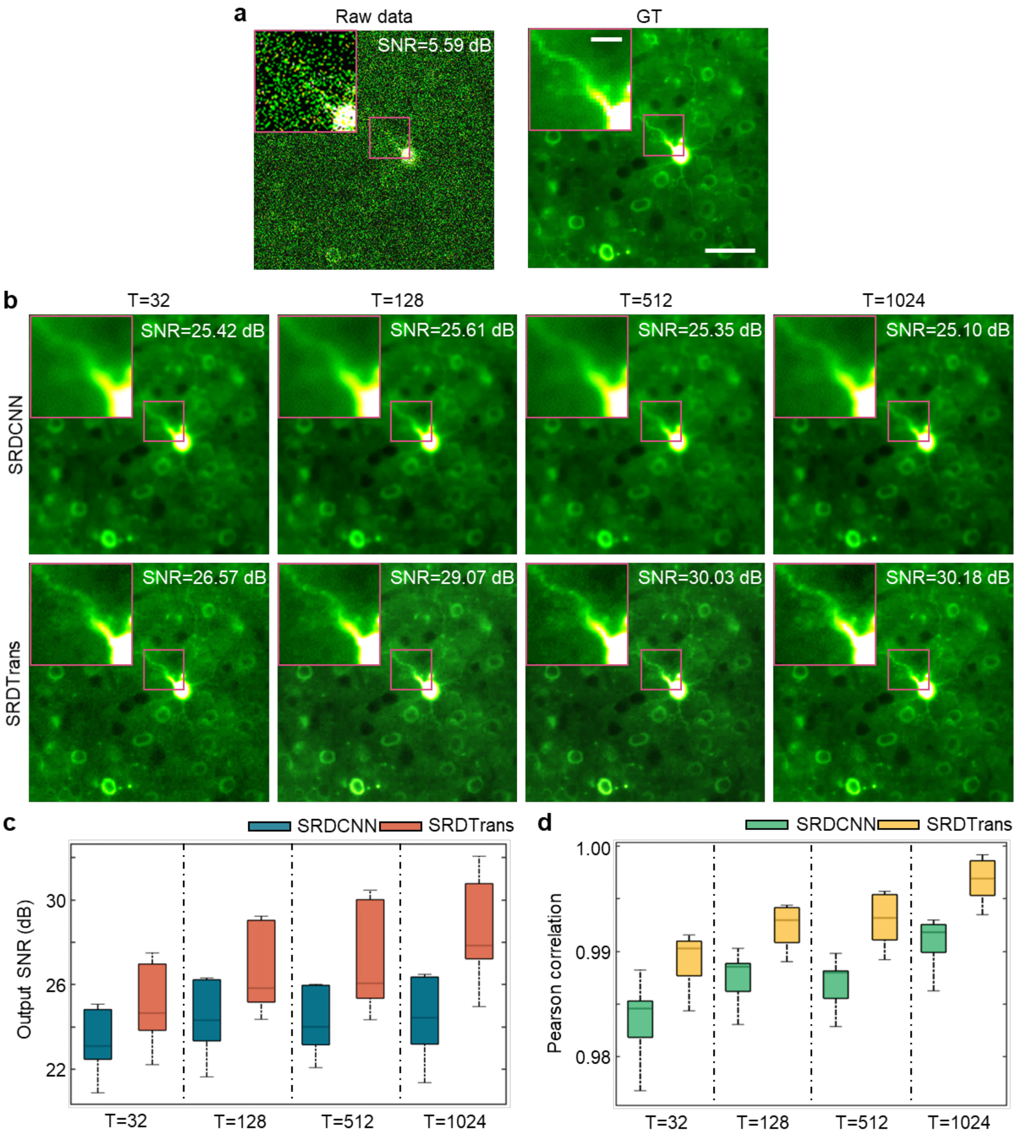


**Supplementary Figure 6**

**Performance of SRDCNN and SRDTrans with different input temporal scales (30-Hz calcium imaging data).**

**a,** A representative frame of simulated calcium imaging data (6000 frames, 30 Hz frame rate, SNR=5.59 dB) and the corresponding ground truth (GT). Scale bar, 40 μm for the whole FOV and 10 μm for magnified views. **b,** SRDCNN and SRDTrans denoised images with different input temporal scales (T). For model training, the input patch size was 128×128×T pixels, where T=32, 128, 512, 1024**. c, d**, Quantitative comparison of SRDCNN and SRDTrans (N=6000). Both SNR (**c**) and Pearson correlation coefficients (**d**) were used as the performance metric. As the input temporal scale increases, more temporal information can be perceived and utilized by SRDTrans, and thus better denoising performance can be obtained and more details can be restored. In contrast, SRDCNN does not show obvious improvement and suffers from blurring.

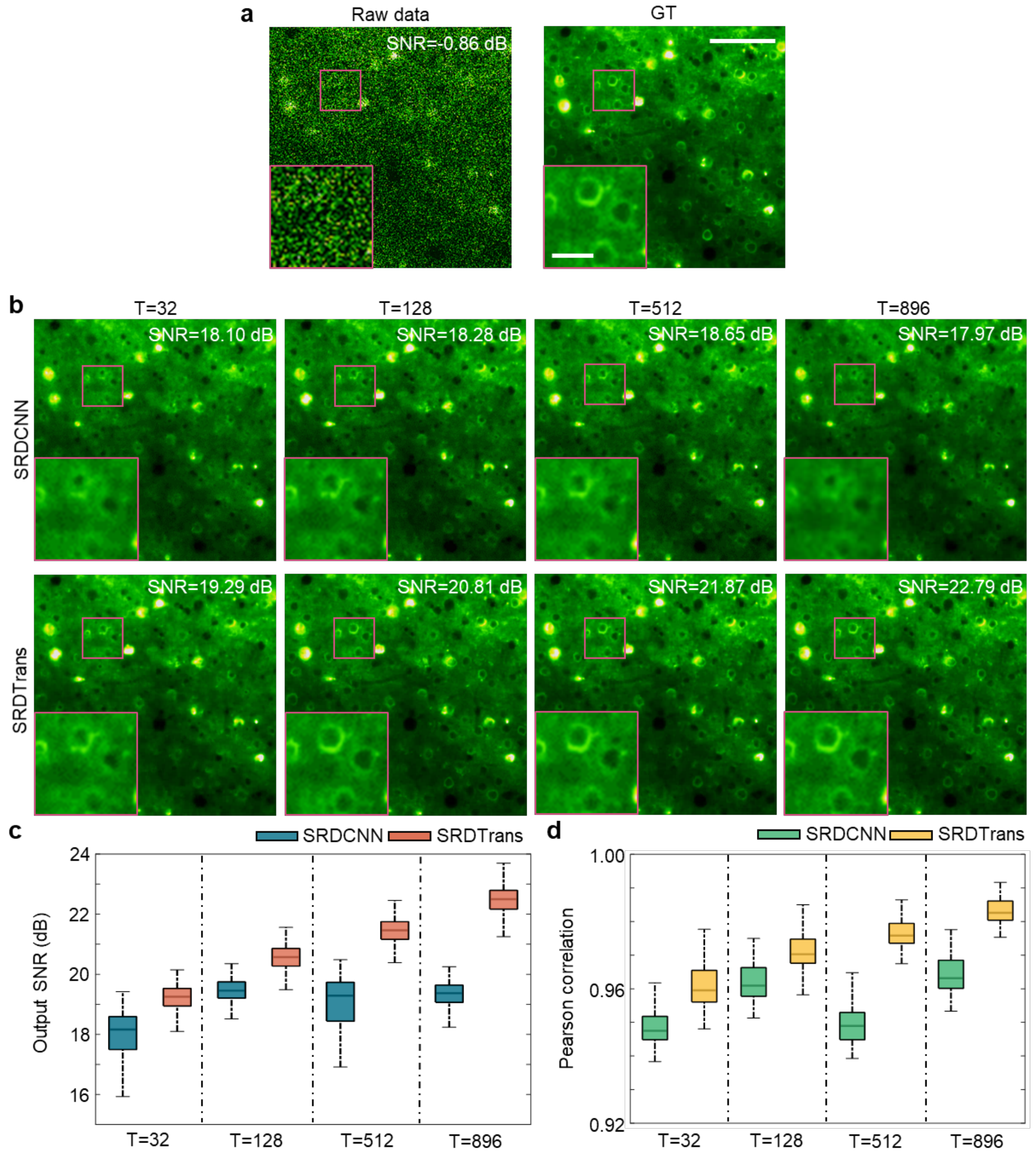


**Supplementary Figure 7**

**Performance of SRDCNN and SRDTrans with different input temporal scales (1-Hz calcium imaging data).**

**a,** A representative frame of simulated calcium imaging data (1000 frames, 1 Hz frame rate, SNR=-0.86 dB) and the corresponding ground truth (GT). Scale bar, 80 μm for the whole FOV and 20 μm for magnified views. **b,** SRDCNN and SRDTrans denoised images with different input temporal scales (T). For model training, the input patch size was 128×128×T pixels, where T=32, 128, 512, 896. **c, d**, Quantitative comparison of SRDCNN and SRDTrans (N=1000). Both SNR (**c**) and Pearson correlation coefficients (**d**) were used as the performance metric. As the input temporal scale increases, more temporal information can be perceived and utilized by SRDTrans, and thus better denoising performance can be obtained and more details can be restored. In contrast, SRDCNN does not show obvious improvement and suffers from blurring.

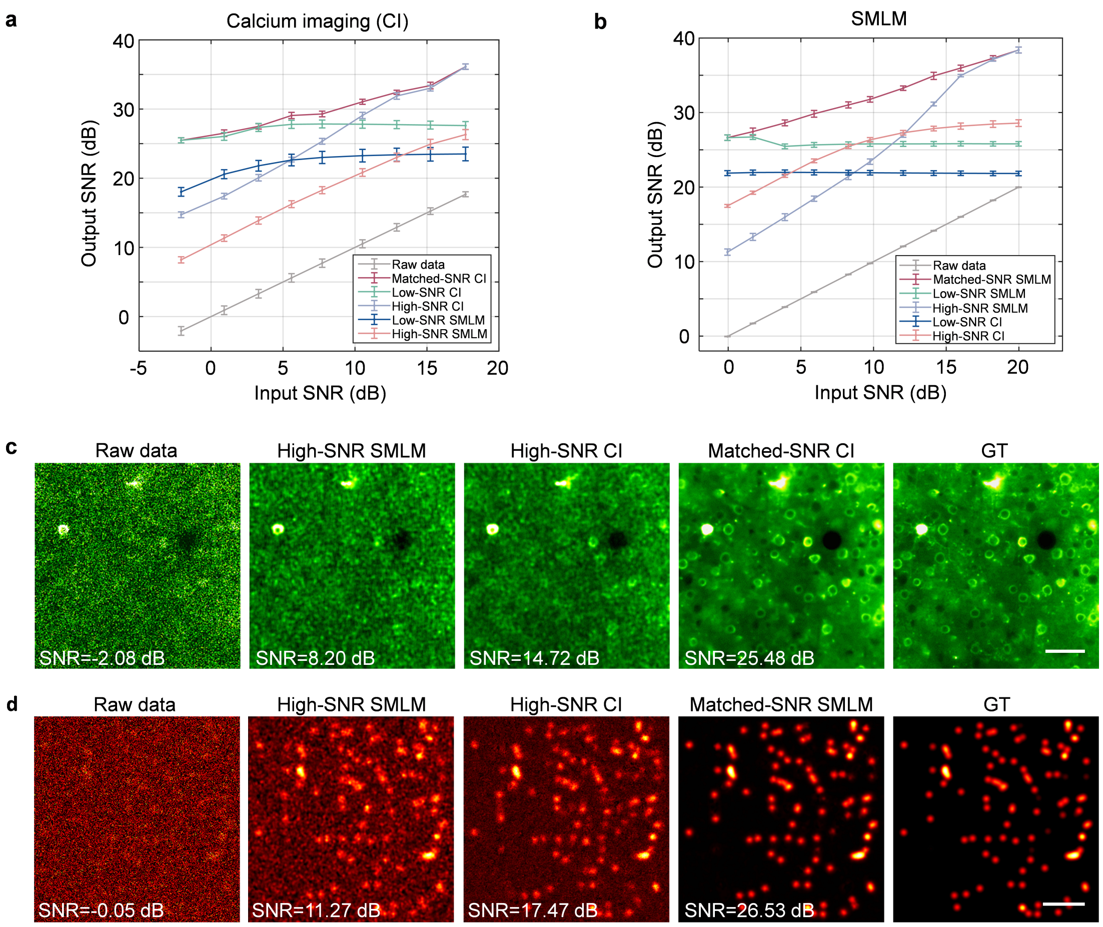


#### Supplementary Figure 8

**Investigating the generalization ability of SRDTrans by cross-dataset and cross-modality validation.**

Denoising performance of models trained on different input SNRs and imaging modalities. **a,** Calcium imaging (CI) data (30 Hz, 6000 frames) of different SNRs were used for test. All values are shown as mean ± s.d. (N=6000). Raw data (gray): raw data without denoising; Matched-SNR CI (dark red): models were trained on CI data with the same SNR as the test data. Low-SNR CI (green): the model was only trained on low-SNR (-2.08 dB) CI data; High-SNR CI (blue grey): the model was only trained on high-SNR (17.86 dB) CI data; Low-SNR SMLM (blue): the model was only trained on low-SNR (-0.05 dB) SMLM data; High-SNR SMLM (light red): the model was only trained on high-SNR (19.98 dB) SMLM data. **b,** Single-molecule localization microscopy (SMLM) data (200 Hz, 24000 frames) of different SNRs were used for test. All values are shown as mean ± s.d. (N=24000). Raw data (gray): raw data without denoising; Matched-SNR SMLM (dark red): models were trained on SMLM data with the same SNR as the test data. Low-SNR SMLM (green): the model was only trained on low-SNR (-2.08 dB) SMLM data; High-SNR SMLM (blue grey): the model was only trained on high-SNR (19.98 dB) SMLM data; Low-SNR CI (blue): the model was only trained on low-SNR (-2.08 dB) CI data; High-SNR CI (light red): the model was only trained on high-SNR (17.86dB) CI data. **c**, Representative images of the results in **a**. A low-SNR (-2.08 dB) CI stack was used to test different models. Scale bar, 50 μm. **d**, Representative images of the results in **b**. A low-SNR (-0.05 dB) SMLM stack was used to test different models. Scale bar, 2 μm.

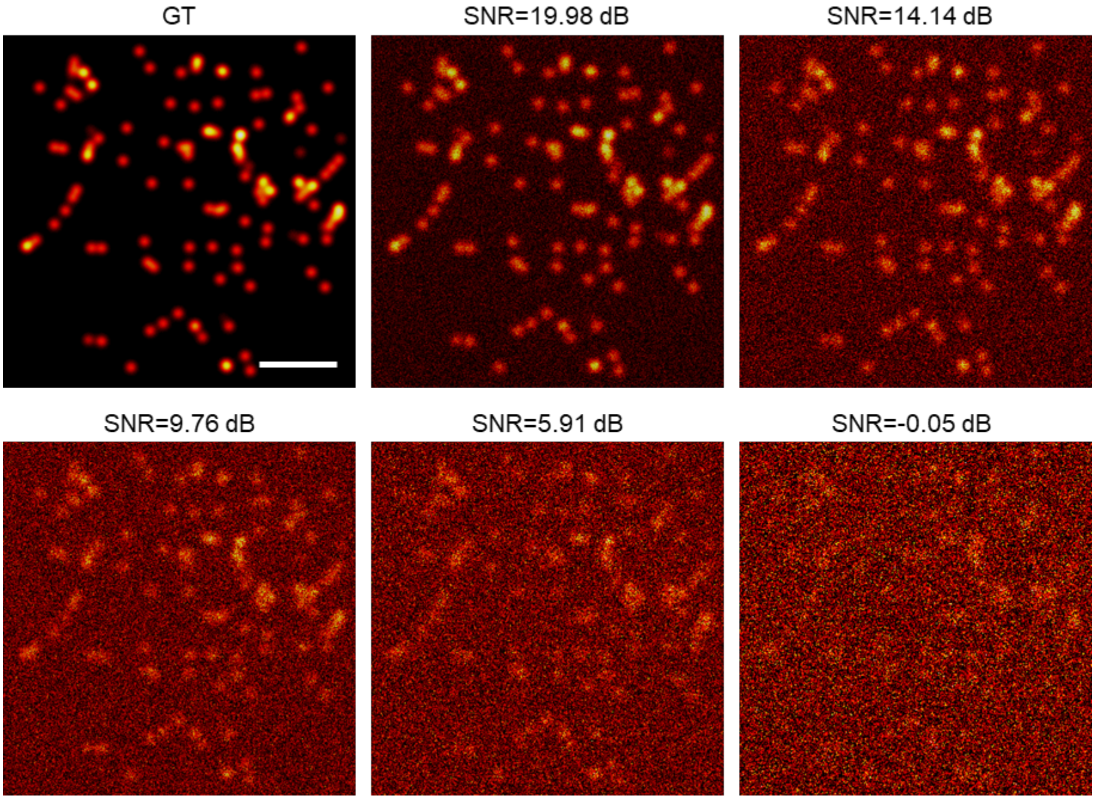


**Supplementary Figure 9**

**Examples of simulated SMLM data with different SNRs.**

We synthesized SMLM data (24000 frames, 200 Hz frame rate) of microtubules with a semi-automated process. The distribution of microtubules was manually extracted from experimentally obtained data released by the ShareLoc.XYZ platform[9] using the JFilament plugin[10]. Next, single-molecule emission images were automatically generated using TestSTORM[11]. Mixed Poisson Gaussian (MPG) noise was further applied to generate authentic low-SNR images. Scale bar, 2 μm.

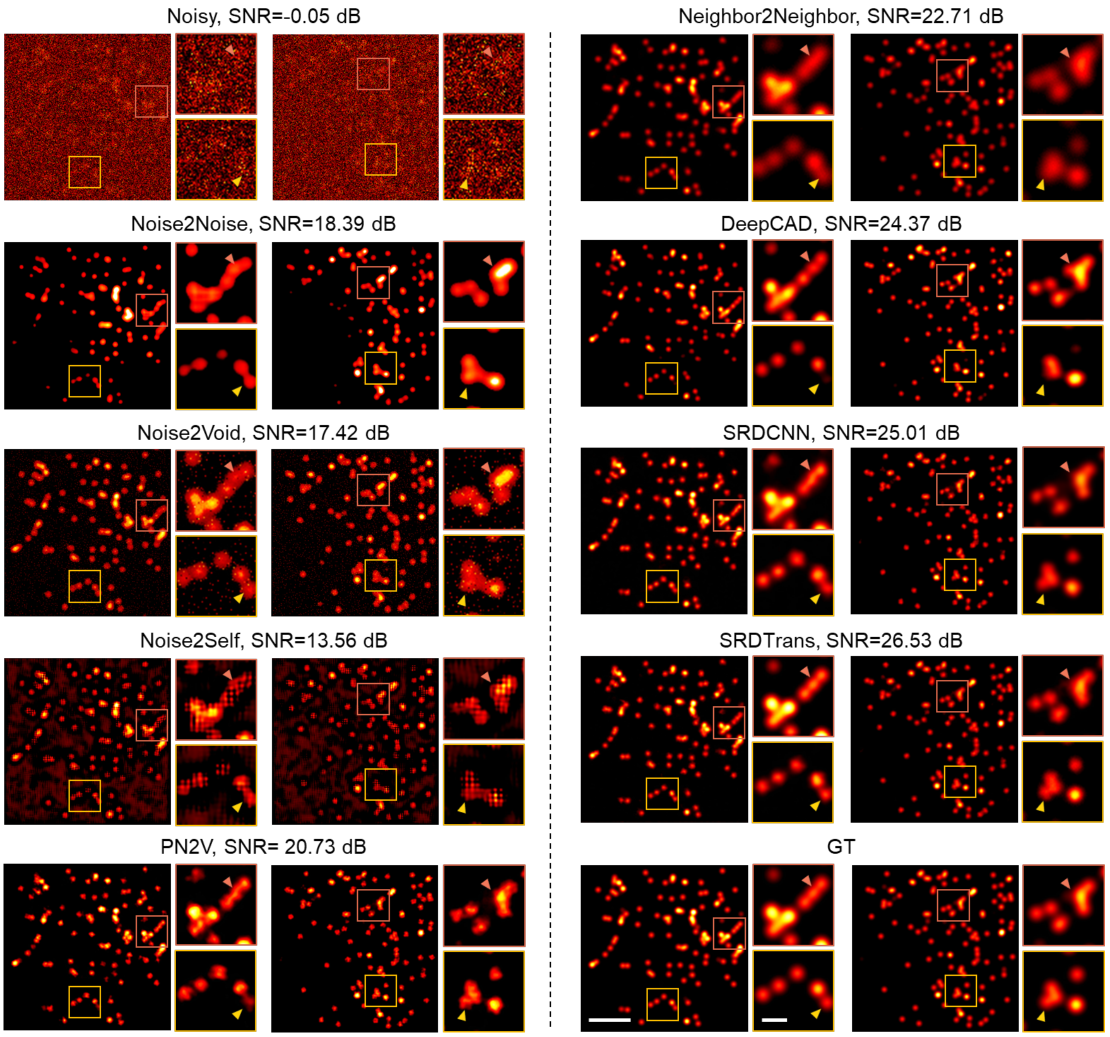


**Supplementary Figure 10**

**Performance of various self-supervised methods.**

Representative results of different denoising methods, including Noise2Noise[12], Noise2Void[1], Noise2Self[2], Probabilistic Noise2Void[13] (PN2V), Neighbor2Neighbor[3], DeepCAD[14,15], and SRDTrans. Simulated SMLM data (24000 frames, 200 Hz frame rate, SNR=-0.05 dB) were used to train specific models for each method. All existing methods were reproduced with the released code of relevant papers and trained with recommended configurations (Supplementary Tab. 4). The noise-free ground truth (GT) is presented in the bottom right panel. As shown in the magnified views, SRDTrans can restore fluorescent molecules with the highest accuracy among all methods. Scale bars, 2 μm for the whole FOV and 0.5 μm for magnified views.

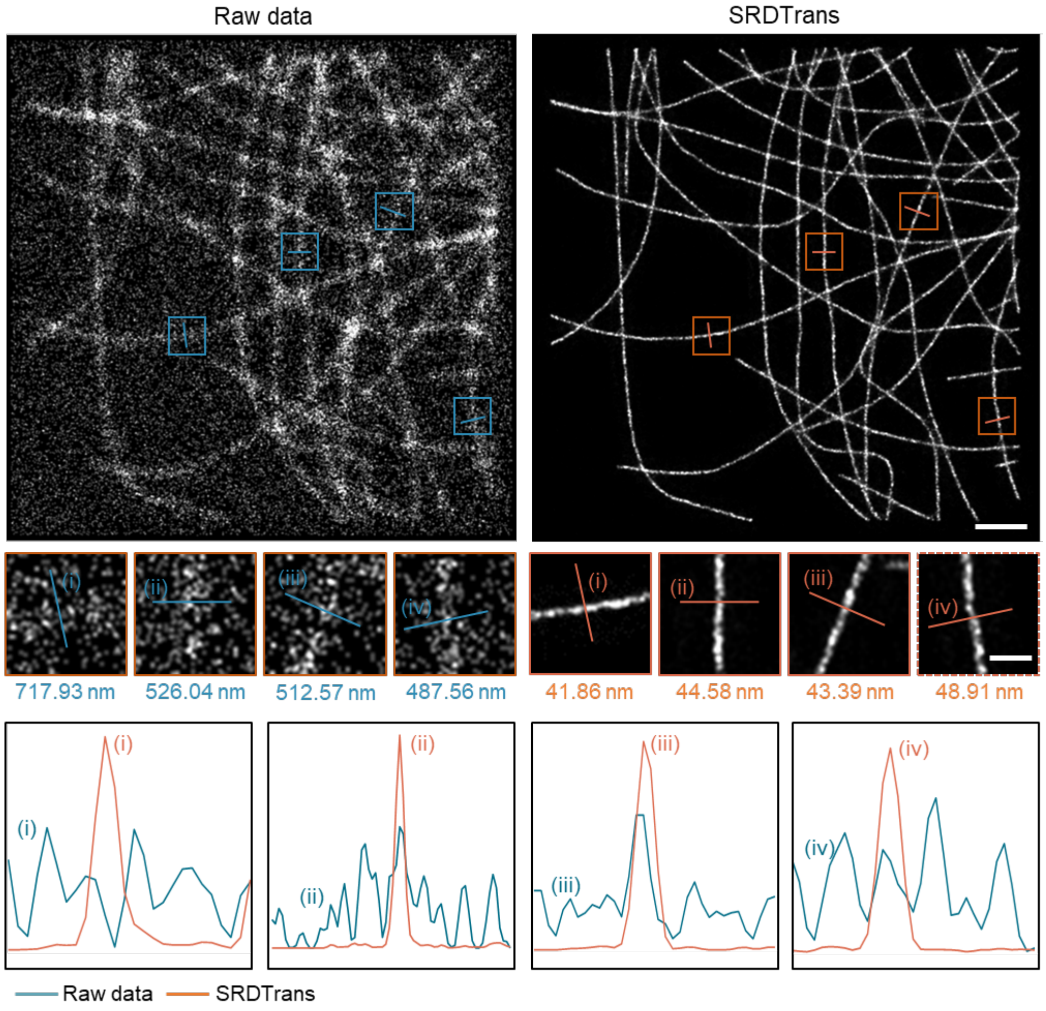


**Supplementary Figure 11**

**Resolution analysis of reconstructed SMLM images.**

Super-resolution images reconstructed by the ThunderSTORM[16] Fiji plugin before (left) and after (right) denoising. Magnified views of the boxed regions show four separate microtubule fragments. Their intensity profiles are plotted in the bottom panel. We applied Gaussian fitting on the intensity profile and measured the full width at half maximum (FWHM, labeled under each magnified view) to reflect the spatial resolution. Disturbed by noise, microtubules in the raw data cannot be well reconstructed and the resolution is severely degraded. In contrast, clear and high-resolution structures can be reconstructed from SRDTrans denoised images. Scale bars, 2 μm for the whole FOV and 0.5 μm for magnified views.

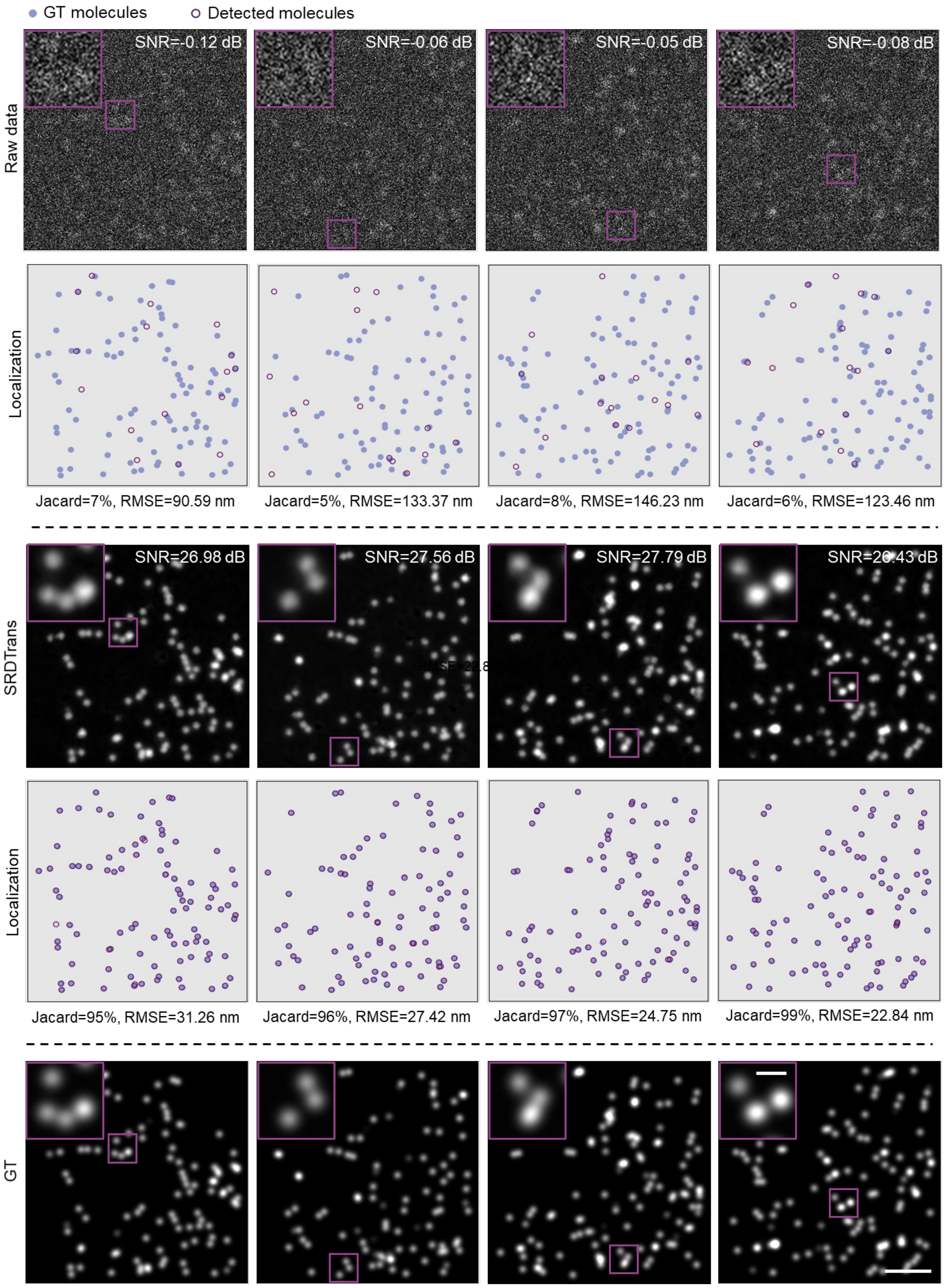
GT molecules
Detected molecules
Raw data
SNR=-0.12 dB
SNR=-0.06 dB
SNR=-0.05 dB
SNR=-0.08 dB
Localization
Jacard=7%, RMSE=90.59 nm
Jacard=5%, RMSE=133.37 nm
Jacard=8%, RMSE=146.23 nm
Jacard=6%, RMSE=123.46 nm
SRDTrans
SNR=26.98 dB
SNR=27.56 dB
SNR=27.79 dB
SNR=26.43 dB
Localization
Jacard=95%, RMSE=31.26 nm
Jacard=96%, RMSE=27.42 nm
Jacard=97%, RMSE=24.75 nm
Jacard=99%, RMSE=22.84 nm
GT

**Supplementary Figure 12**

**Localization accuracy before and after denoising.**

Four representative frames of the raw SMLM data (top panel), SRDTrans denoised data (middle panel), and ground truth (GT, bottom panel). The SNR of each image is labeled. The ThunderSTORM[16] Fiji plugin was used for single-molecule localization, and the localization result is shown under each image. Blue spots indicate GT molecules and purple circles indicate detected molecules. Scale bars, 2 μm for the whole FOV and 0.5 μm for magnified views. The Jaccard index and root mean squared error (RMSE) were calculated to quantitatively evaluate the localization accuracy. Disturbed by noise, fewer molecules were localized from the raw noisy data, and the localization accuracy was low. SRDTrans can increase the number of detectable molecules and localization accuracy.

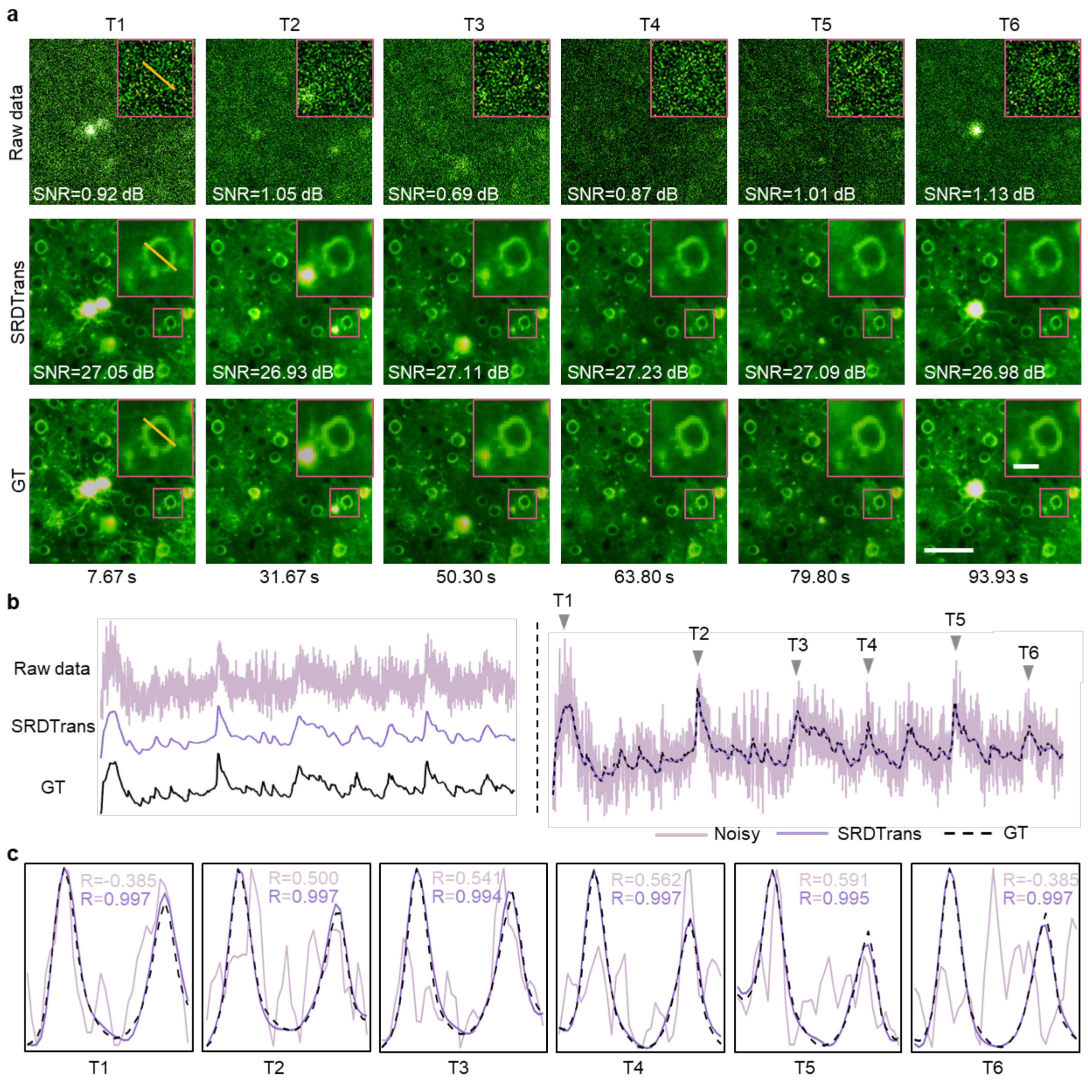


#### Supplementary Figure 13

**Spatiotemporal analysis of denoised calcium imaging data.**

Simulated calcium imaging image data (6000 frames, 30 Hz frame rate) was used in this experiment. **a,** Representative frames of the raw data, SRDTrans enhanced data, and corresponding ground truth (GT). Six different moments are presented and the SNR of each image is annotated. Scale bars, 50 μm for the whole FOV and 10 μm for magnified views. **b,** The calcium trace of a cytoplasmic pixel before and after denoising. **c,** Pixle intensity along the yellow line indicated in **a**. Pearson correlation coefficients with respect to GT were calculated to quantify the similarity. All Pearson correlation coefficients were improved after SRDTrans denoising.

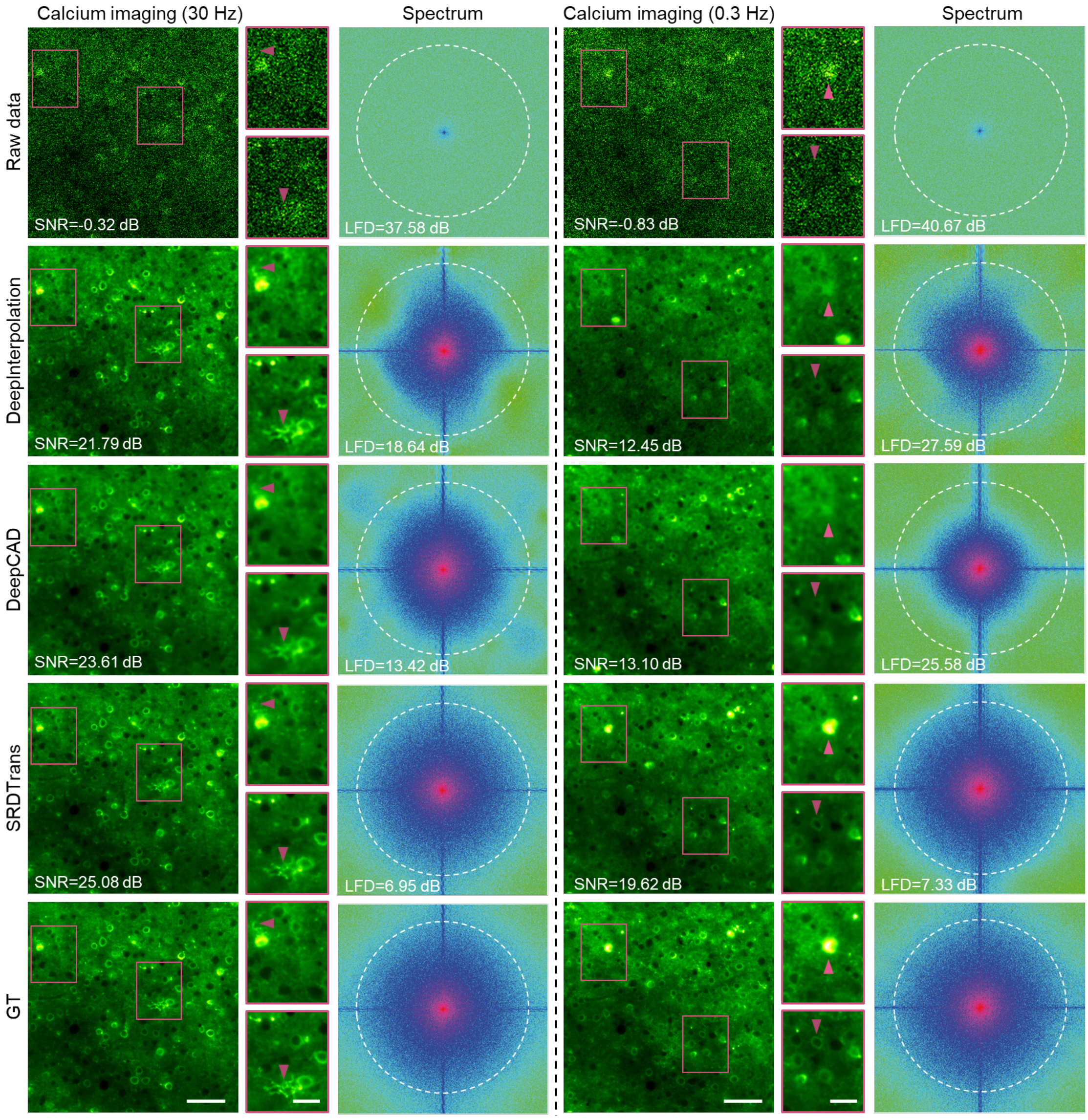


**Supplementary Figure 14**

**Frequency domain analysis of different denoising methods.**

To demonstrate the superiority of SRDTrans in recovering high-frequency information, we visualized the spatial image and Fourier spectrum of the results produced by SRDTrans, DeepInterpolation[17], and DeepCAD. The raw noisy images and ground-truth (GT) images are also presented. Simulated calcium imaging data (1000 frames) with a frame rate of 30 Hz (left column) and 0.3 Hz (right column) were used for comparison. The arrowheads indicate some salient calcium events. The Fourier spectrum shows that DeepCAD and DeepInterpolation would lose high-frequency information while SRDTrans could nearly recover the entire spectrum. SNR and logarithmic frequency distance (LFD) were used for quantification in the spatial domain and the frequency domain, respectively. Scale bars, 50 μm for the whole FOV and 20 μm for magnified views.

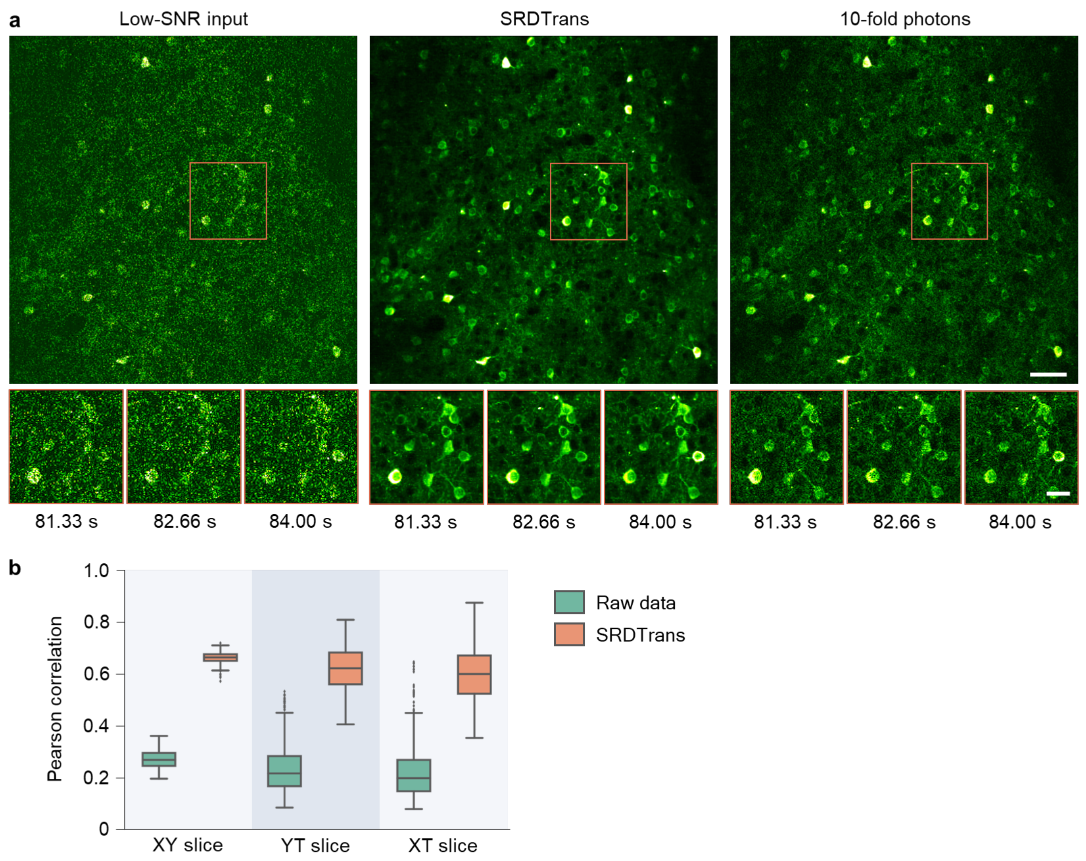


#### Supplementary Figure 15

**Denoising experimentally obtained calcium imaging data with synchronized high-SNR (10-fold photons) reference.**

**a**, Calcium imaging data before and after denoising. The same frame is shown for the original low-SNR data (left), SRDTrans denoised image (middle), and synchronized high-SNR image with tenfold fluorescence photons (right). The data was captured by a custom two-photon microscope on the mouse cortex expressing genetically encoded GCaMP6f calcium indicator[15]. The frame rate of the original data (30 Hz) was down-sampled to 3 Hz by extracting one frame every ten frames. Magnified views of the red-boxed region show the calcium dynamics of a group of neurons. Scale bars, 50 µm for the whole FOV and 20 µm for magnified views. **b**, Box plots showing image correlations along the three dimensions (x-y-t) before and after denoising. The high-SNR data with tenfold fluorescence photons were used as the reference for correlation computing; XY slice, N = 600; YT slice, N = 490; x-t slice, N = 490.

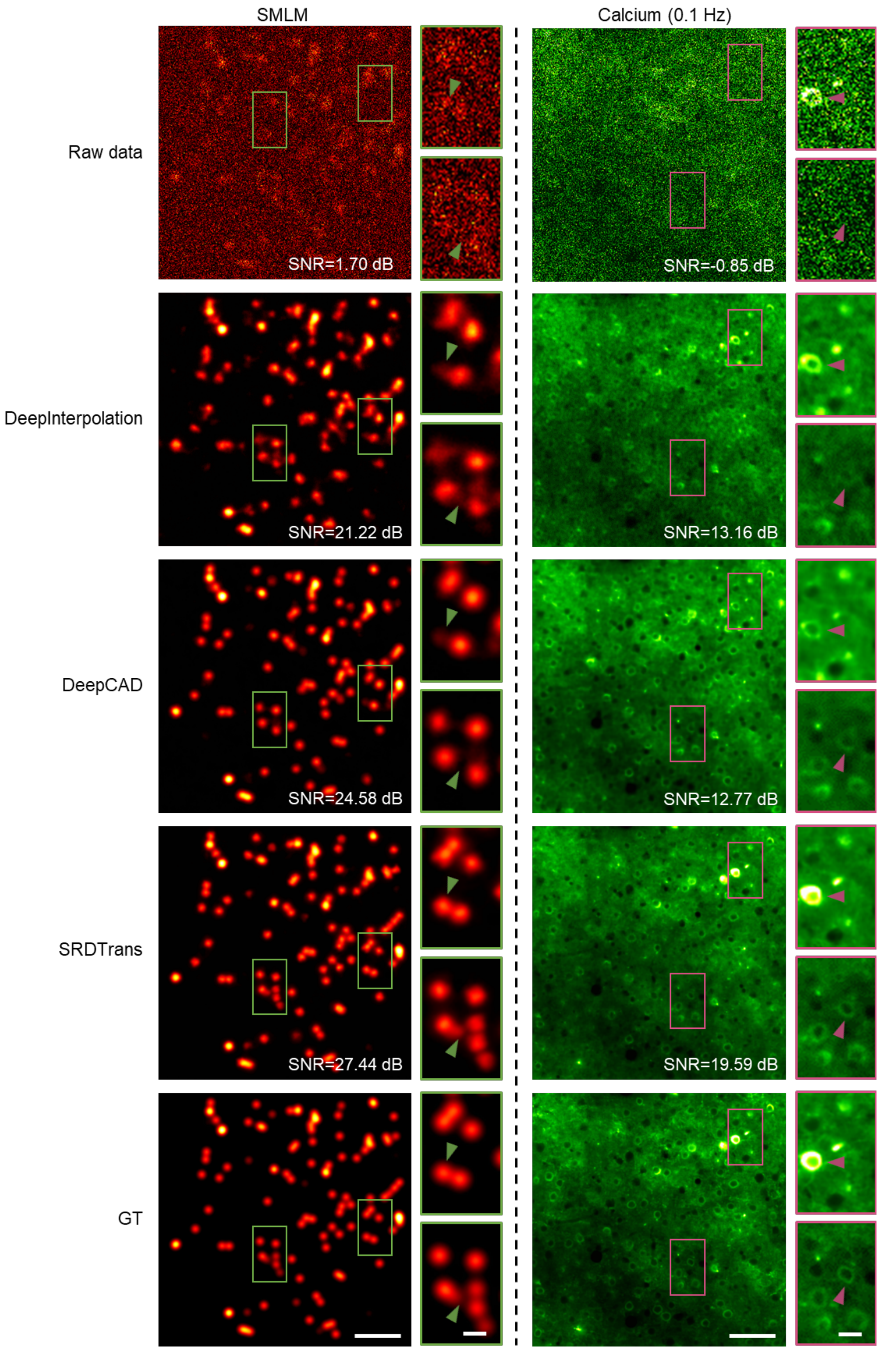


**Supplementary Figure 16**

**Comparing the performance of DeepInterpolation, DeepCAD, and SRDTrans.**

We trained a specific model for each method and each imaging modality with simulated SMLM data (24000 frames, 200 Hz frame rate, SNR=1.70 dB, left column) and calcium imaging data (1000 frames, 0.1 Hz frame rate, SNR=-0.85 dB, right column). The results of different methods are demonstrated here. Magnified views of the boxed regions are shown alongside each image. The arrowheads indicate some salient structures. Scale bar, 2 μm for the whole FOV and 0.5 μm for magnified views (SMLM); 50 μm for the whole FOV and 10 μm for magnified views (calcium imaging).

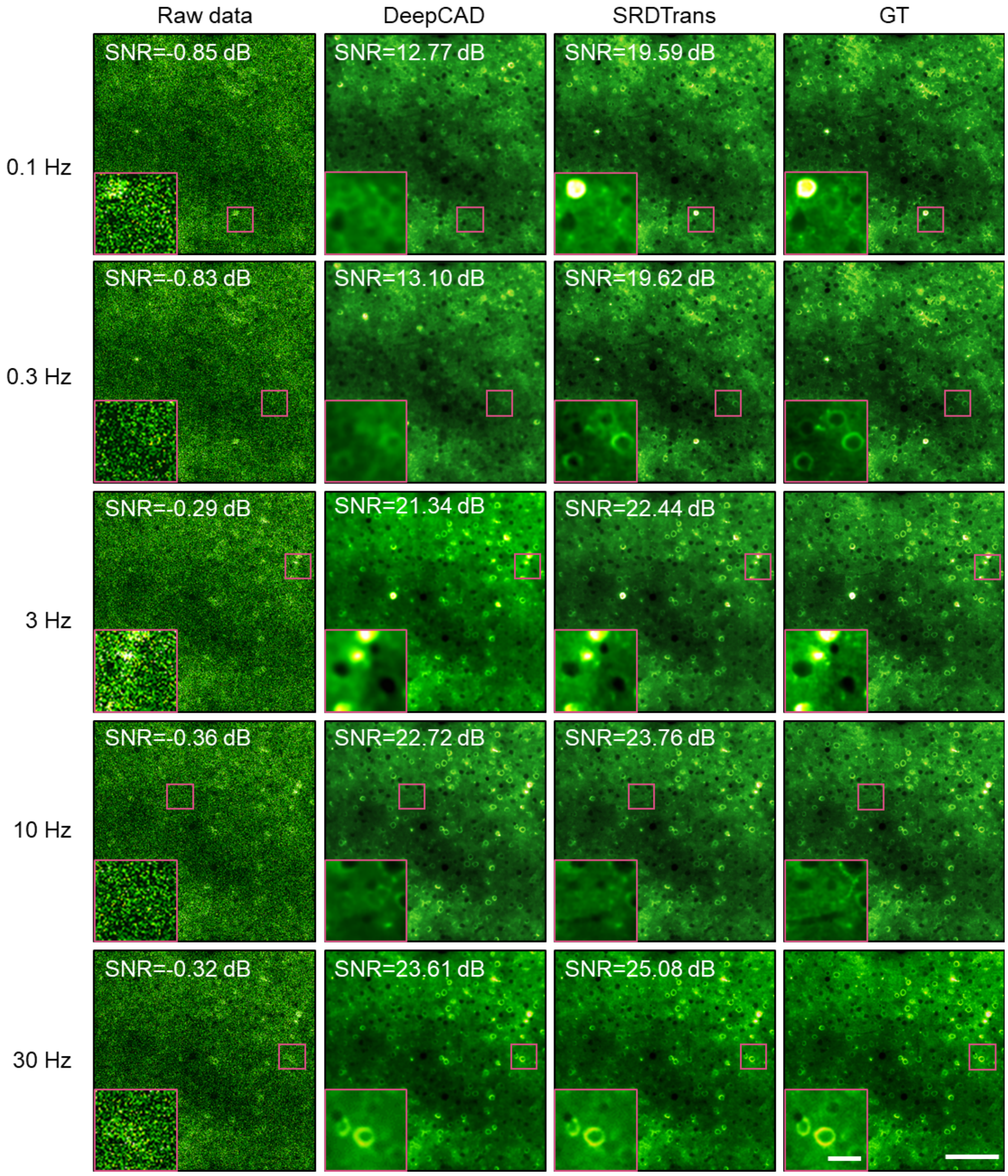


**Supplementary Figure 17**

**Performance of DeepCAD and SRDTrans at different imaging speeds.**

We compared the denoising performance of DeepCAD and SRDTrans at different imaging speeds. Simulated calcium imaging data (1000 frames) were used to train specific models for each method and each frame rate. As calcium transients change rapidly, a lower frame rate means that the similarity between two adjacent frames is low. Since DeepCAD was designed to utilize temporal redundancy, its performance drops rapidly as the frame rate decreases. In contrast, SRDTrans combines spatial redundancy sampling and a transformer architecture. It can achieve better and more stable performance than DeepCAD at various imaging speeds. Scale bars, 50 μm for the whole FOV and 10 μm for magnified views.

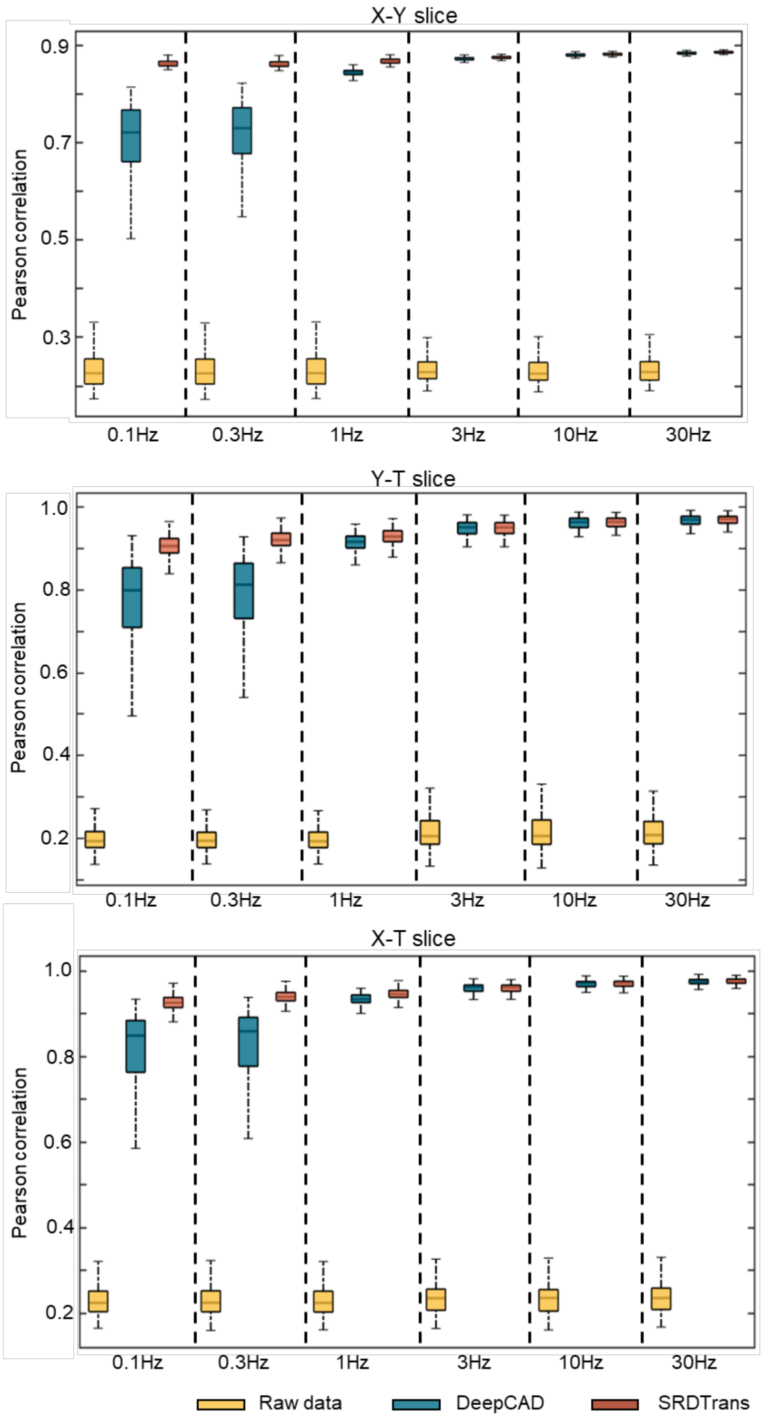


**Supplementary Figure 18**

**Pearson correlations at different imaging speeds.**

Box plots showing image correlations along the three dimensions (x-y-t) before and after denoising. The ground-truth images were used as the reference for correlation computing. Simulated calcium imaging data of different frame rates were used in this experiment. Top, X-Y slice, N =1,000. Middle, Y-T slice, N = 490. Bottom, X-T slice, N = 490.

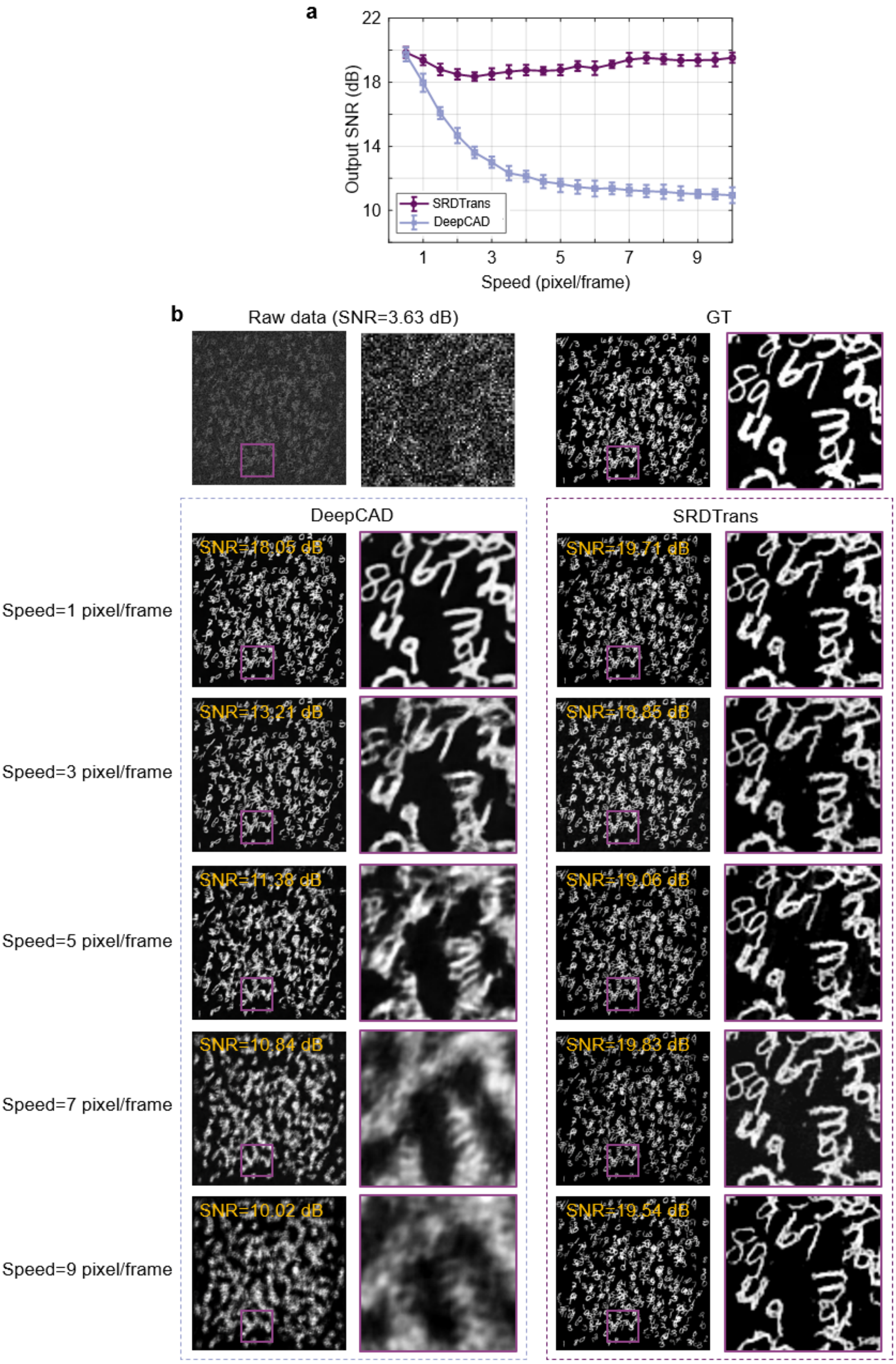
a
Output SNR (dB)
22
18
14
10
1
3
5
7
9
Speed (pixel/frame)
SRDTrans
DeepCAD
b
Raw data (SNR=3.63 dB)
GT
DeepCAD
SRDTrans
Speed=1 pixel/frame
Speed=3 pixel/frame
Speed=5 pixel/frame
Speed=7 pixel/frame
Speed=9 pixel/frame
SNR=18.05 dB
SNR=19.71 dB
SNR=13.21 dB
SNR=18.85 dB
SNR=11.38 dB
SNR=19.06 dB
SNR=10.84 dB
SNR=19.83 dB
SNR=10.02 dB
SNR=19.54 dB

**Supplementary Figure 19**

**Comparing DeepCAD and SRDTrans on moving objects with different moving speeds.**

We synthesized moving objects with different moving speeds to verify the robustness of SRDTrans on samples undergoing fast dynamics. **a**, Quantitative comparison of DeepCAD and SRDTrans on moving objects with different moving speeds (N=5000 frames for each moving speed). The lines connect mean values, and the error bars represent standard deviation (N=5000). **b**, Representative results of DeepCAD and SRDTrans at the speed=1,3,5,7,9 pixel/frame. The raw noisy data and corresponding ground truth (GT) are presented in the top panel. Each individual digit was randomly sampled from the MNIST[18] dataset. See Supplementary Video 4 for the video.

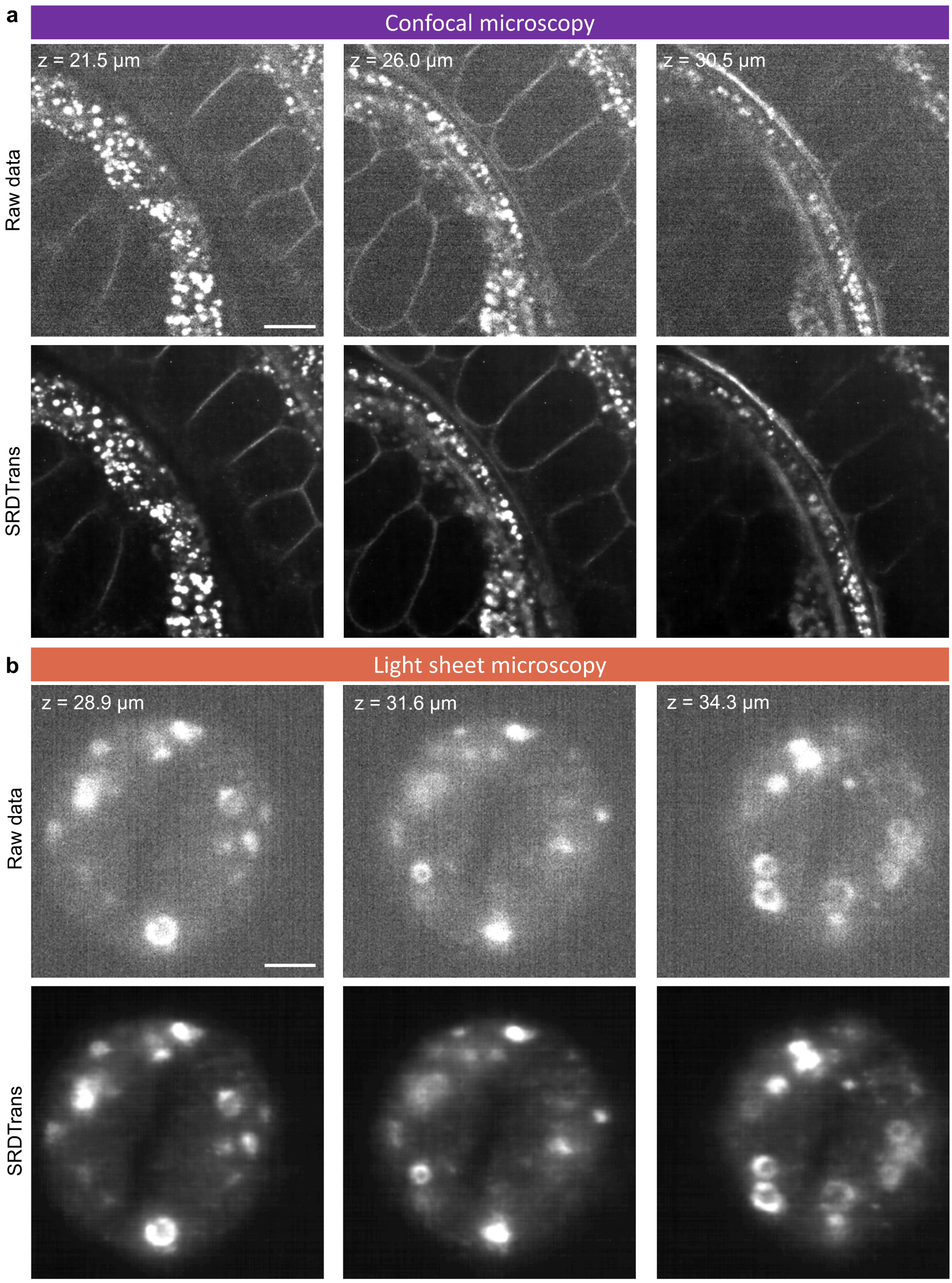
a
Confocal microscopy
z = 21.5 µm
z = 26.0 µm
z = 30.5 µm
Raw data
SRDTrans
b
Light sheet microscopy
z = 28.9 µm
z = 31.6 µm
z = 34.3 µm
Raw data
SRDTrans

**Supplementary Figure 20**

**The applicability of SRDTrans on confocal microscopy and light sheet microscopy.**

**a**, Denoising performance of SRDTrans on confocal microscopy. Representative images of three axial layers are presented. The data was acquired by axially scanning a mRuby3 labeled *C. elegans* using a commercial confocal microscope (SpinSR, Olympus). Scale bar, 20 µm. **b**, Denoising performance of SRDTrans on light sheet microscopy. Representative images of three axial layers are presented. The data was acquired by axially scanning a Lamp1-Halo labeled HeLa cell using a lattice light-sheet microscope[19]. The fluorescence signals are from lysosomes. Scale bar, 5 µm. The model for each imaging modality was trained and tested on a single volumetric (xy-z) stack without any reliance on temporal information.

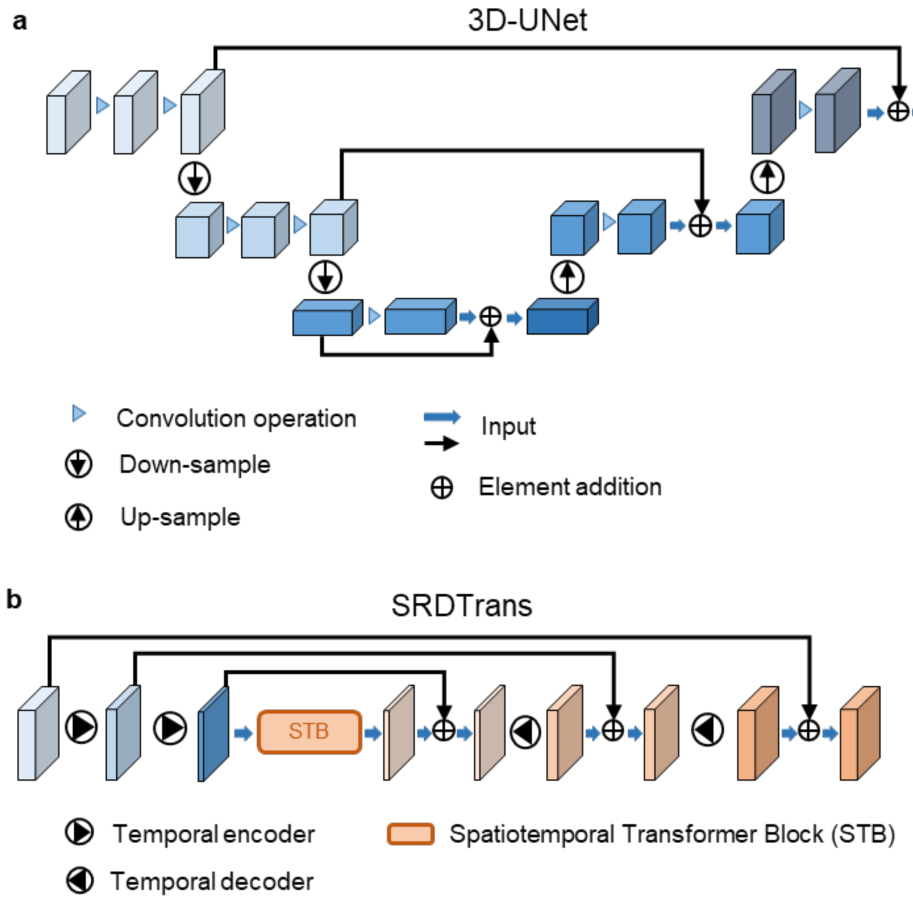


**Supplementary Figure 21**

**The mechanism of feature extraction in 3D-UNet and SRDTrans.**

**a**, A typical 3D-UNet employs a U-shaped structure to encode the input image into multi-scale features and then decode it to the original scale[20]. The key unit of the encoding and decoding module is the convolution layers, and the receptive field is limited by the size of convolution kernels. Moreover, the up-sampling and down-sampling operations lower the spatial resolution of the features, resulting in limited ability in high-frequency information extraction. **b**, SRDTrans does not apply any spatial up-sampling and down-sampling to allow more high-frequency information to pass. The spatiotemporal transformer block (STB) endows SRDTrans with the ability to extract global information both in space and time.

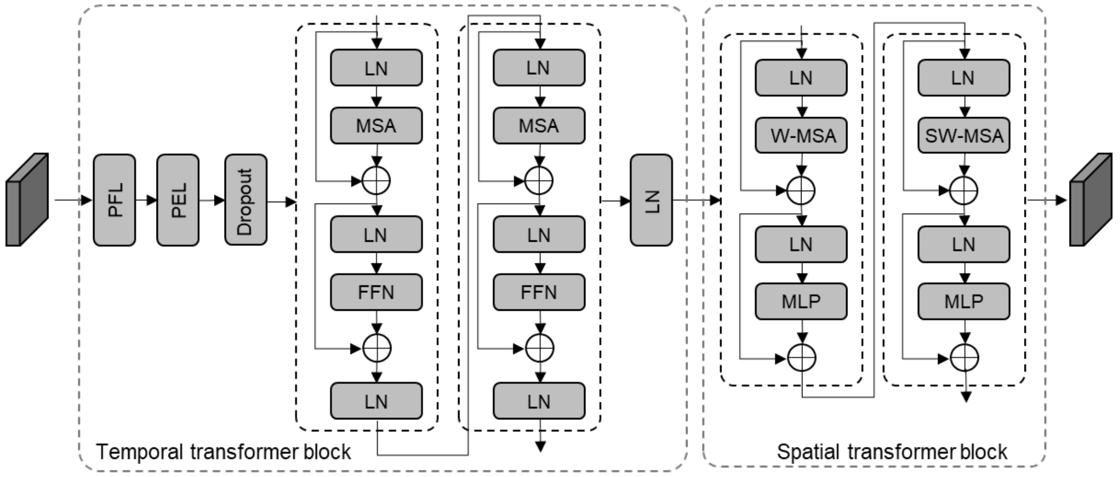


PFL: Patch flattening layer
PEL: Position embedding layer
LN: Layer normalization
MSA: Multi-head self-attention
FFN: Feed-forward network
W-MSA: Window-based multi-head self-attention
SW-MSA: Shifted window-based multi-head self-attention
MLP: Multi-layer perceptron

**Supplementary Figure 22**

**Architecture of the spatiotemporal transformer block (STB).**

The proposed STB is composed of a temporal transformer block and a spatial transformer block. The temporal transformer block comprises a patch flattening layer (PFL), a position embedding layer (PEL), a dropout layer, two self-attention blocks, and a layer normalization (LN). Each self-attention block implements the standard multi-head self-attention (MSA) mechanism to capture long-range temporal dependencies. The spatial transformer block comprises a Swin Transformer block[21]. W-MSA and SW-MSA are multi-head self-attention modules with regular and shifted windowing configurations, respectively. The hierarchical architecture is beneficial for learning fine-grained spatial features.

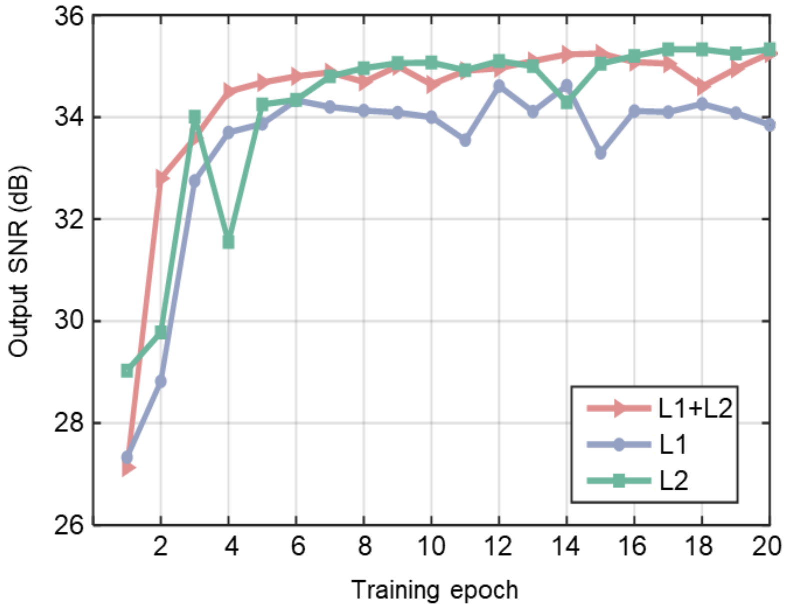


**Supplementary Figure 23**

**Convergence curves of different loss functions.**

The network was trained on simulated calcium imaging data (6000 frames, 30 Hz frame rate, SNR=9.69 dB) for 20 epochs with different loss functions (L1, L2, L1+L2). The performance (average SNR) after each epoch was tested. All parameters remained the same except the loss function.

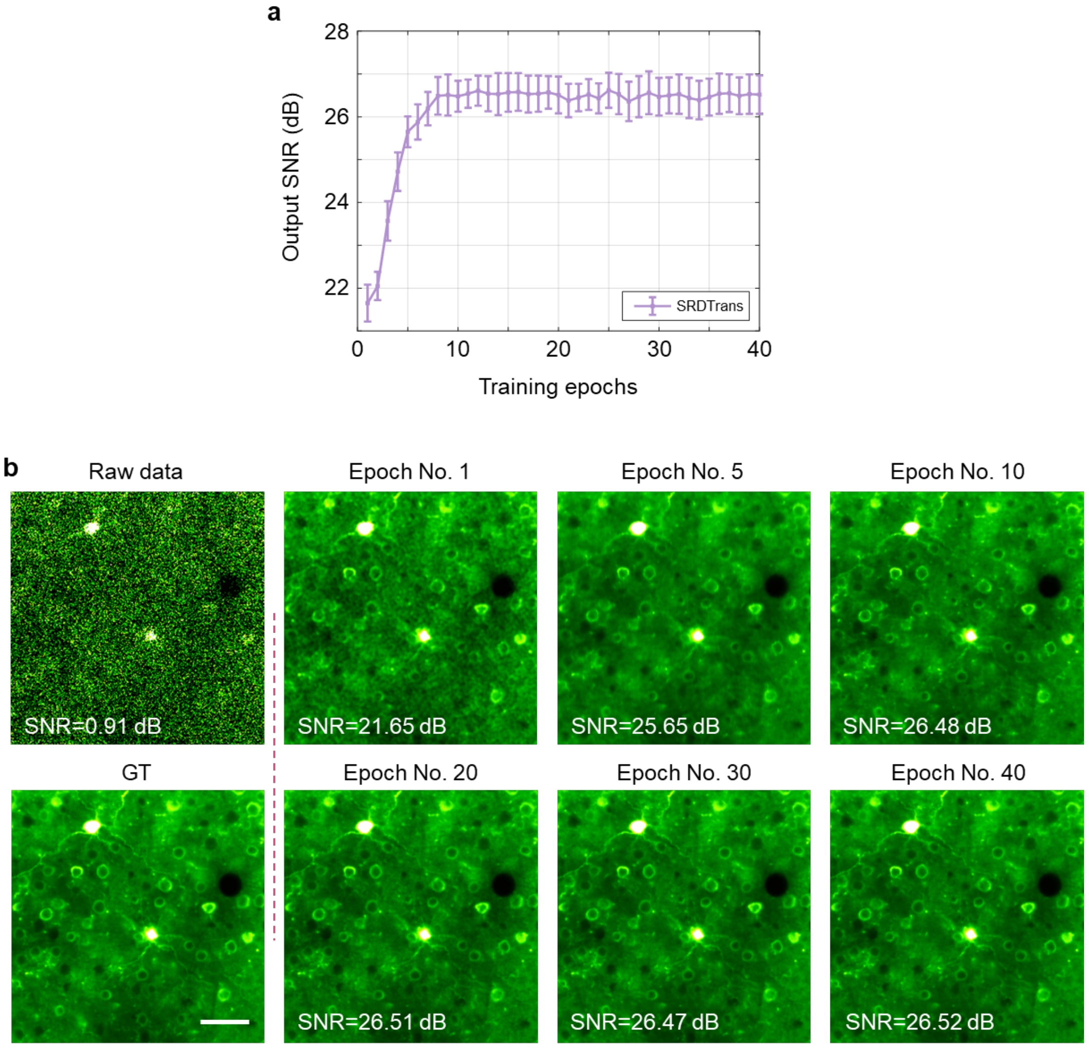


**Supplementary Figure 24**

**Training stability of SRDTrans.**

Simulated calcium imaging data (6000 frames, 30 Hz frame rate, SNR=0.91 dB) were used in this experiment for quantitative evaluation. The network was trained for 40 epochs and the model of each epoch was tested. **a**, Denoising performance of SRDTrans with the increase of training epoch. All values are shown as mean ± s.d. (N=6000). **b**, Example denoised images of different epochs. The raw noisy data and corresponding ground truth (GT) are presented in the leftmost column. SRDTrans shows strong training stability and resistance to overfitting. Scale bar, 50 μm.

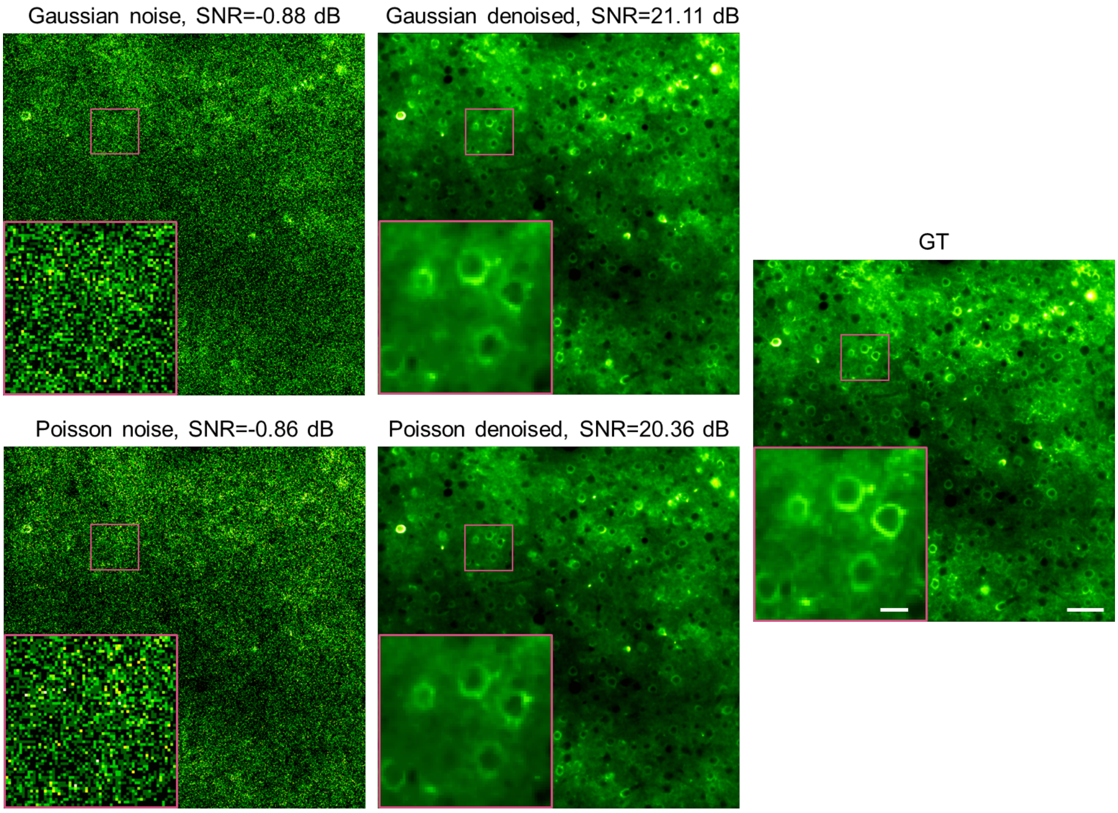


**Supplementary Figure 25**

**Comparing the denoising performance of SRDTrans on Gaussian and Poisson noise.**

Simulated calcium imaging data (1000 frames, 30 Hz frame rate) containing different forms of noise (Gaussian noise and Poisson noise) but with the same SNR were generated. SRDTrans were used to denoise these data with the same training and testing hyperparameters. The average SNR is labeled above each image and the corresponding ground truth (GT) is presented in the rightmost column. Scale bars, 50 μm for the whole FOVs and 10 μm for magnified views.

### Supplementary Table

#### Supplementary Table 1

##### Model complexity of CNN and SRDTrans.

We compared the number of parameters, inference time, and denoising performance (quantified by SNR) of CNN and SRDTrans with different feature channels. We used 3D-UNet as the representative CNN architecture in this experiment. Simulated calcium imaging data (N=6000 frames, 30 Hz frame rate, SNR=10.52 dB) were used as the training and testing dataset. For each method, we trained the network for 20 epochs and selected the optimal model for inference. The output SNRs (mean ± s.d.) are summarized in the bottom line of the table. The GPU used for training and inference is Nvidia GeForce 2080Ti.

| **The number of feature channels** | | **8** | **16** | **32** | **64** | **128** | **256** |
|---|---|---|---|---|---|---|---|
| **Parameters (M)** | CNN | 0.26 | 1.02 | 4.08 | 16.32 | 65.26 | 261.01 |
| | SRDTrans | 0.62 | 0.68 | 0.82 | 1.11 | 1.79 | 3.55 |
| **Inference Time (ms)** | CNN | **8.12** | **14.38** | **32.78** | 109.23 | 387.13 | 1538.53 |
| | SRDTrans | 47.83 | 50.93 | 54.12 | **59.15** | **72.03** | **104.56** |
| **Output SNR (dB)** | CNN | 27.43 ±0.29 | 28.18 ±0.37 | 28.33 ±0.21 | 28.01 ±0.23 | 28.22 ±0.39 | 28.29 ±0.27 |
| | SRDTrans | **29.53** ±0.34 | **30.51** ±0.22 | **30.66** ±0.16 | **30.79** ±0.27 | **30.91** ±0.21 | **31.04** ±0.35 |

**Supplementary Table 2**

**Model complexity and denoising performance of different transformer-style networks.**

For model complexity analysis, we compared SRDTrans with several commonly used transformer-style networks applicable for time-lapsed imaging data, including Video Swin Transformer (VST)[22], nnFormer[23], UNETR[24], and Swin UNETR[25]. All methods were reproduced with the released code of relevant papers and trained with default configurations. The training and testing dataset was simulated calcium imaging data (N=6000 frames, 30 Hz frame rate, SNR=10.52 dB). During training, the patch size was 128×128×128 pixels. Since the VST was designed for short-sequence videos, its complexity on long-sequence is too high to operate on our device, so we only provide model parameters for reference. All models were trained with the Nvidia GeForce 2080Ti GPU.

| | **Parameters ↓ (M)** | **Inference time ↓ (ms)** | **Running memory ↓ (GB)** | **Output SNR ↑ (dB)** |
|---|---|---|---|---|
| VST | 224.68 | -* | -* | -* |
| nnFormer | 71.90 | 238.41 | 15.24 | 28.33 |
| UNETR | 93.01 | 267.56 | 19.38 | 26.62 |
| Swin UNETR | 62.20 | 98.76 | 8.76 | 28.54 |
| **SRDTrans** | **1.79** | **72.03** | **5.22** | **30.91** |

* It cannot be obtained because the model complexity exceeds the limits of the device.

**Supplementary Table 3**

**Quantitative comparison of SRDCNN, DeepCAD, and SRDTrans at different imaging speeds.**

We compared SRDTrans with SRDCNN and DeepCAD on simulated calcium imaging data at different imaging speeds. Simulated calcium imaging data (N=1000 frames) sampled at 0.1 Hz, 0.3 Hz, 1 Hz, 3 Hz, 10 Hz, and 30 Hz were used as the training and testing datasets. These methods combining different network architectures and sampling strategies. For SRDCNN, we trained a 3D-UNet with the proposed spatial redundancy sampling. DeepCAD is based on temporal redundancy and used temporally adjacent frames to train a 3D-UNet. SRDTrans combines spatial redundancy sampling and a lightweight 3D-Transfomer network. The output SNRs (mean ± s.d.) are summarized in this table and representative images are shown in Supplementary Fig. 17. These results quantitatively show that (1) spatial redundancy sampling is superior to temporal redundancy sampling when the imaging speed is low. (2) The backbone 3D-Transfomer network of SRDTrans can provide better denoising performance than 3D-UNet.

| **Frame rate** | | **0.1 Hz** | **0.3 Hz** | **1 Hz** | **3 Hz** | **10 Hz** | **30 Hz** |
|---|---|---|---|---|---|---|---|
| **SRDCNN** | 3D-UNet / Spatial redundancy | 17.66 ±0.43 | 18.04 ±0.56 | 18.67 ±0.64 | 20.61 ±0.33 | 21.64 ±0.28 | 22.41 ±0.37 |
| **DeepCAD** | 3D-UNet / Temporal redundancy | 12.77 ±2.01 | 13.10 ±1.61 | 18.44 ±0.87 | 21.34 ±0.25 | 22.72 ±0.22 | 23.61 ±0.22 |
| **SRDTrans** | 3D-Transfomer / Spatial redundancy | **19.59** ±0.37 | **19.62** ±0.31 | **20.83** ±0.36 | **22.44** ±0.30 | **23.76** ±0.30 | **25.08** ±0.23 |

**Supplementary Table 4**

**Parameters for the SMLM experiments.**

All the SMLM experiments in this paper were conducted on a commercial microscope Nikon N-STORM. The detailed parameters are listed as follows.

| **Physiological parameters** | | **Imaging parameters** | |
|---|---|---|---|
| Cell type | BSC-1 | FOV | 11 × 11 μm |
| Labeling | Cy5 | Pixel size | 43nm |
| Organelles | Microtubules | Image size | 256×256 pixels |
| Laser source | | Detection NA | 1.4 |
| Excitation wavelength | 640 nm | Magnification | 150 |
| Excitation power | 80mW | Number frames | 60000 or 120000 |
| Beam concentration | 4× | Exposure time | 5ms |
| Activation wavelength | 405 nm | Frame rate | 200 Hz |
| Activation power | 20mW | | |

**Supplementary Table 5**

**Training configurations of different self-supervised denoising methods.**

We used the released code of relevant papers and follow their recommended configurations. The patch size of all methods was the same to ensure that they had access to the same information during training. The detailed training settings for each method are listed follows.

| Method | Patch size | Network | BS | lr | Loss | Epoch |
|---|---|---|---|---|---|---|
| **Noise2Noise** | 128×128 | U-Net | 16 | $1e^{-3}$ | $L_1$ | 300 |
| **Noise2Void** | 128×128 | U-Net | 128 | $4e^{-4}$ | $L_1$ | 100 |
| **Noise2Self** | 128×128 | U-Net | 64 | $1e^{-3}$ | $L_1$ | 50 |
| **Neighbor2Neighbor** | 128×128 | U-Net | 4 | $3e^{-4}$ | $L_2+L_{reg}$ | 100 |
| **DeepInterpolation** | 128×128×128 | 3D-UNet | 1 | $1e^{-4}$ | $L_1$ | 50 |
| **DeepCAD** | 128×128×128 | 3D-UNet | 1 | $5e^{-5}$ | $L_{1+}\ L_2$ | 20 |
| **SRDTrans** | 128×128×128 | 3D-Transformer | 1 | $1e^{-4}$ | $L_{1+}\ L_2$ | 20 |

a. BS: batch size.
b. lr: learning rate

### Supplementary Notes

#### 1. Proof of spatial redundancy sampling scheme.

We first prove the rationality of training the denoising network with two randomly sampled sub-stacks $y_1$ and $y_2$ from a single image stack $y$. We assume that $y_1$ and $y_2$ are two independent noisy stacks conditioned on the noise-free stack $x$. Since the noise of each pixel is independent, the sub-stack pair ($y_1$, $y_2$) can be considered as two independent samples of the same underlying pattern[3,26], we have:

$$
\begin{aligned}
\mathrm{E}_{y_1|x}\left|\mathrm{F}_\theta(y_1)-x\right|_2 &= \mathrm{E}_{y_1,y_2|x}\left|\mathrm{F}_\theta(y_1)-y_2+y_2-x\right|_2 \\
&= \mathrm{E}_{y_1,y_2|x}\left|\mathrm{F}_\theta(y_1)-y_2\right|_2+\mathrm{E}_{y_2|x}\left|y_2-x\right|_2+2\mathrm{E}_{y_1,y_2|x}(\mathrm{F}_\theta(y_1)-y_2)^\tau(y_2-x) \\
&= \mathrm{E}_{y_1,y_2|x}\left|\mathrm{F}_\theta(y_1)-y_2\right|_2+\sigma_{y_2}^2+2\mathrm{E}_{y_1,y_2|x}(\mathrm{F}_\theta(y_1)-x+x-y_2)^\tau(y_2-x) \\
&= \mathrm{E}_{y_1,y_2|x}\left|\mathrm{F}_\theta(y_1)-y_2\right|_2+\sigma_{y_2}^2+2\mathrm{E}_{y_1,y_2|x}(\mathrm{F}_\theta(y_1)-x)^\tau(y_2-x) \\
&\quad +2\mathrm{E}_{y_2|x}(x-y_2)^\tau(y_2-x) \\
&= \mathrm{E}_{y_1,y_2|x}\left|\mathrm{F}_\theta(y_1)-y_2\right|_2-\sigma_{y_2}^2+2\mathrm{E}_{y_1,y_2|x}(\mathrm{F}_\theta(y_1)-x)^\tau(y_2-x)
\end{aligned}
$$

where $\sigma_{y2}$ is the variance of $y_2$. And we assume that there exists an $\varepsilon \neq 0$ such that $\mathrm{E}(y_1) = x$ and $\mathrm{E}(y_2) = x + \varepsilon$. Since $y_1$ and $y_2$ are independent under the condition of given $x$, the further derivation can be obtained as:

$$
\begin{aligned}
\mathrm{E}_{y_1|x}\left|\mathrm{F}_\theta(y_1)-x\right|_2 &= \mathrm{E}_{y_1,y_2|x}\left|\mathrm{F}_\theta(y_1)-y_2\right|_2-\sigma_{y_2}^2+2\mathrm{E}_{y_1|x}(\mathrm{F}_\theta(y_1)-x)^\tau\mathrm{E}_{y_2|x}(y_2-x) \\
&= \mathrm{E}_{y_1,y_2|x}\left|\mathrm{F}_\theta(y_1)-y_2\right|_2-\sigma_{y_2}^2+2\varepsilon\mathrm{E}_{y_1|x}(\mathrm{F}_\theta(y_1)-x).
\end{aligned}
$$

Then we have:

$$
\mathrm{E}_{x,y_1}\left|\mathrm{F}_\theta(y_1)-x\right|_2 = \mathrm{E}_{x,y_1,y_2}\left|\mathrm{F}_\theta(y_1)-y_2\right|_2-\sigma_{y_2}^2+2\varepsilon\mathrm{E}_{x,y_1}(\mathrm{F}_\theta(y_1)-x),
$$

If $y_1$ and $y_2$ are similar enough, $\varepsilon$ is close to 0. Then we can use the network output trained with noisy sub-stack ($y_1$, $y_2$) as a reasonable approximation to the supervised learning with ground-truth $x$.

#### 2. The spectral bias of CNN

When fitting the objective function, CNNs usually tend to give priority to fitting low-frequency information and finally to high-frequency information[8]. The spectral bias phenomenon caused by the F-Principle[6] makes it challenging to fit high-frequency components inside the signal. Here we explain the necessity of correcting the spectral bias in CNNs in image restoration following some proven theoretical frameworks and proofs in previous work[7,8].

We first clarify the theoretical understanding of the **F-principle**. The Fourier transformation and its inverse transformation are defined as follows:

$$F(w)=\int_{-\infty}^{+\infty} f(x)^{-wx}x,$$
$$f(x)=\frac{1}{2\pi}\int_{-\infty}^{+\infty} F(w)^{wx}w.$$

We define the activation function as $\sigma(kx+b)$, and the Fourier transform of $\sigma$ is:

$$\sigma(w)=\frac{2\pi}{k}e^{\frac{bw}{k}}\frac{1}{e^{-\frac{\pi w}{2k}}-e^{\frac{\pi w}{2k}}}.$$

The amplitude deviation between the DNN output at the frequency $w$ and the objective function $f(x)$ is defined as,

$$\begin{aligned}D(w)&=H(w)-F(w)\\&=\sum_{j=1}^{m}\frac{2\pi a}{k}e^{\frac{bw}{k}}\frac{1}{e^{-\frac{\pi w}{2k}}-e^{\frac{\pi w}{2k}}}-F(w),\end{aligned}$$

and $D(w)$ can be expressed as a combination of magnitude $A(w)$ and phase $\phi(w)$: $D(w)=A(w)^{\phi(w)}$. The loss at frequency $w$ is $L(w)=\frac{1}{2}\left|D(w)\right|^2$, so the total loss function is defined as:

$$L=\int_{-\infty}^{+\infty} L(w)w=\int_{-\infty}^{+\infty}\frac{1}{2}(h(x)-f(x))^2x$$

The partial derivatives of $L(w)$ with respect to parameters $\{a,k,b\}$ are:

$$\frac{\partial L(w)}{\partial a}=\frac{2\pi}{k}\sin(\frac{bw}{k}-\phi(w))E_0,$$
$$\frac{\partial L(w)}{\partial k}=[\sin(\frac{bw}{k}-\phi(w))(\frac{\pi^2 aw}{k^3}E_1-\frac{2\pi a}{k^2})-\frac{2\pi abw}{k^3}\cos(\frac{bw}{k}-\phi(w))]E_0,$$
$$\frac{\partial L(w)}{\partial b}=\frac{2\pi abw}{k^2}\cos(\frac{bw}{k}-\phi(w))]E_0,$$

where

$$E_0=\frac{\operatorname{sgn}(k)A(w)}{e^{\frac{\pi w}{2k}}-e^{-\frac{\pi w}{2k}}},$$
$$E_1=\frac{e^{\frac{\pi w}{2k}}+e^{-\frac{\pi w}{2k}}}{e^{\frac{\pi w}{2k}}-e^{-\frac{\pi w}{2k}}}.$$

If the delta is reduced relative to the parameter $\theta$, for example: $\frac{\partial L}{\partial \theta}=\int_{-\infty}^{+\infty}\frac{\partial L(w)}{\partial \theta}w$.The effect on the total amount at frequency k can be expressed as:

$$\left|\frac{\partial L(w)}{\partial \theta}\right|\approx A(w)e^{-\left|\frac{\pi w}{2k}\right|}N(\theta,w),\quad \theta\triangleq\left\{k,b,a\right\},$$

When the component $H(w)$ at frequency $w$ is not close enough to $F(w)$, and the $e^{-\frac{\pi w}{2k}}$ will dominate the gradient component. We consider an activation function $\sigma(x)=\tanh(x)$ or $\sigma(x)=\max(0,x)$ of the CNN with a hidden layer, for any frequency $w_1$ and $w_2$ satisfying $|F(w_1)|>0,\ |F(w_2)|>0$ and $|w_2|>|w_1|>0$, there are constants $c$ and $C$ such that for small enough δ, we get:

$$\frac{\mu\left(\left\{k:\left|\frac{\partial L(w_1)}{\partial \theta}\right|>\left|\frac{\partial L(w_2)}{\partial \theta}\right|\right\}\cap B_\delta\right)}{\mu(B_\delta)}\geq 1-Ce^{-\frac{c}{\delta}},$$

where $B_\delta \subset \mathbb{R}^m$ is a ball with radius $\delta$ centered at the origin and $\mu(\cdot)$ is the Lebesgue measure. The F-principle theorem states that for any two non-convergent frequencies with small weights, low-frequency gradients outperform high-frequency gradients. According to Parseval's theorem, the MSE loss in the spatial domain is equivalent to the L2 loss in the Fourier domain. Preferential fitting of CNNs to low frequencies by itself results in synthetic differences in the frequency domain, which is also the main reason for the over-smoothing structures in CNN restoration results.
